# Probiotic phage steering of disease suppressive rhizosphere microbiome

**DOI:** 10.64898/2026.01.02.697320

**Authors:** Keming Yang, Xiaofang Wang, Jingxuan Li, Xinyu Tang, Yilin He, Yike Tang, Shuo Wang, Rujiao Hou, Yangchun Xu, Qirong Shen, Ville-Petri Friman, Zhong Wei

## Abstract

Current phage therapy approaches predominantly focus on targeting pathogenic bacteria, overlooking the vast potential of phages that interact with beneficial, plant growth-promoting (probiotic) bacteria. Here we tested if a single phage targeting a probiotic *Stenotrophomonas maltophilia* bacterium can steer the rhizosphere microbiome disease suppressiveness against *Ralstonia solanacearum* phytopathogen. We find that *S. maltophilia* quickly evolves resistance to its phage via mutations in *ssb* and *TonB* genes and by upregulating anti-phage defense systems. Crucially, evolution of phage resistance reprograms the bacterial transcriptome and metabolome, leading to unexpectedly enhanced antimicrobial activity against the *R. solanacearum*. Furthermore, exposing *S. maltophilia* to phage in the tomato rhizosphere increases the microbiota-wide disease suppressiveness by stabilizing bacterial diversity and facilitating pathogen suppression by other resident species. Our findings suggest that probiotic-specific phages could be used as ecological and evolutionary engineers to steer the rhizosphere microbiome disease suppressiveness through activation of antagonistic bacterial interactions.

## INTRODUCTION

Bacterial viruses - phages - are the most abundant and diverse microbial group on Earth. They play key role in regulating bacterial community diversity, composition and functioning and can indirectly affect the health of associated eukaryotic hosts through effects on host-associated microbiomes^1^. Due to high specificity, phages have traditionally been used to target pathogenic bacterial species in both clinical and agricultural settings^2^. A recent study conducted in the tomato rhizosphere demonstrated that phages targeting plant growth-promoting bacteria can indirectly benefit the growth of *Ralstonia solanacearum* pathogen by reducing the strength of bacterial competition and associated pathogen suppression^3^. Moreover, phages targeting *R. solanacearum* has been shown to indirectly increase the diversity of tomato rhizosphere microbiota by reducing the pathogen densities and vacating niche space for commensal and plant growth-promoting bacteria^4,5^. However, very little is known how phages interact with commensal or beneficial bacteria in eukaryote-associated microbiomes and how this could indirectly affect the pathogen suppression.

In addition to ecological mechanisms, such as bacterial density regulation and species sorting^3,5,6^, phage selection could alter the functioning of plant growth-promoting, probioic bacteria through genetic correlations between phage resistance and other traits^7^. For example, mutations in phage receptor genes (*e.g.*, outer membrane protein modifications) that block phage adsorption are often costly due to antagonistic pleiotropy as these genes are also vital for bacterial growth and other functions such as attachment^8^ or acquisition of iron^9^. However, also synergistic pleiotropy has been observed, where phage resistance correlates positively with other bacterial traits such as antibiotic resistance^10^. For example, a subset population of *E. coli* can gain both phage and tetracycline resistance under phage selection through mutations mapped in or near the surface-exposed loops of TolC^11^. Phages have also been reported to activate bacterial secretion of toxic molecules^6,12,13^ or siderophores^14^, and for example, phage infecting *Brevibacillus laterosporus* bacterium boosted the suppression of pathogenic *Paenibacillus larvae* via increased production of bacterial toxins^15^. It remains however unclear how phage selection affects the activity of plant growth-promoting bacteria, and if these changes can cascade through trophic networks, affecting the functioning of surrounding plant rhizosphere microbiota.

Herein, we used laboratory and tomato rhizosphere microbiome^3^ experiments to study how the evolution of phage resistance of probiotic *Stenotrophomonas maltophilia* bacterium^16^ affects its interaction with a phytopathogenic *Ralstonia solanacearum* bacterium, which causes huge agricultural and economic losses by infecting hundreds of agronomically important crops^17^. By combining genome resequencing with transcriptomics and metabolomics, we identified changes in genes affecting *S. maltophilia* phage resistance and antimicrobial activity and validated the inhibitory activity of identified key metabolites that were secreted by *S. maltophilia* due to phage resistance evolution. Greenhouse experiments were used to validate that phage-resistant *S. maltophilia* mutants show increased pathogen suppression *in planta* and that selection by *S. maltophilia*-specific phage could enrich other rhizosphere bacterial taxa that also exerted antimicrobial activity against *R. solanacearum*. Together, our results show that a phage, which targets a single plant growth-promoting probiotic bacterial taxa, could be used to increase/steer rhizosphere microbiota-wide disease suppressiveness by promoting antagonistic metabolic interactions.

## RESULTS

### Phage selection increases *S. maltophilia* phage resistance and disease suppressiveness through positive trait correlations

To study how phage selection might change the activity of probiotic bacteria, we focused on previously characterized probiotic *S. maltophilia* bacterial strain (YL-STE-01^3^) and its specialist phage, YL-STE-01P^3^, which was verified to be incapable of infecting any of the 88 taxonomically representative bacterial strains obtained from the tomato rhizosphere, including *R. solanacearum* (Fig. S1a-c). To experimentally determine whether *S. maltophilia* can evolve resistance to its phage and if the phage resistance leads to correlated responses with *R. solanacearum* pathogen suppression, we first co-evolved *S. maltophilia* with its phage for two days under controlled laboratory conditions. The densities of *S. maltophilia* populations were clearly suppressed in the presence of phage but not entirely eradicated, indicative of evolution of phage resistance (Fig. 1a). To validate these population level observations at the strain level, we isolated a total of 184 single *S. maltophilia* colonies from phage treatment replicates after 48h of co-culturing and directly examined evolution of phage resistance as the relative difference in bacterial biomass in the absence and presence of phage (‘relative phage resistance’). A total of 95.65% (176/184) of colonies showed significant increase in phage resistance compared to ancestral *S. maltophilia* strain (Unpaired t-test, *P* < 0.05, Fig. 1b, Supplementary data S1) and based on this data, colonies were classified into ‘phage-resistant’ (relative phage resistance > 5%, n = 72) and ‘phage-sensitive’ (relative phage resistance < -5%, n = 28) groups. Phage resistance incurred significant but small growth cost in the absence of phage relative to ‘phage-sensitive’ colonies (6.83% reduction in biomass at 24h time point; Wilcoxon-test, *P* < 0.001).

**Figure 1.**
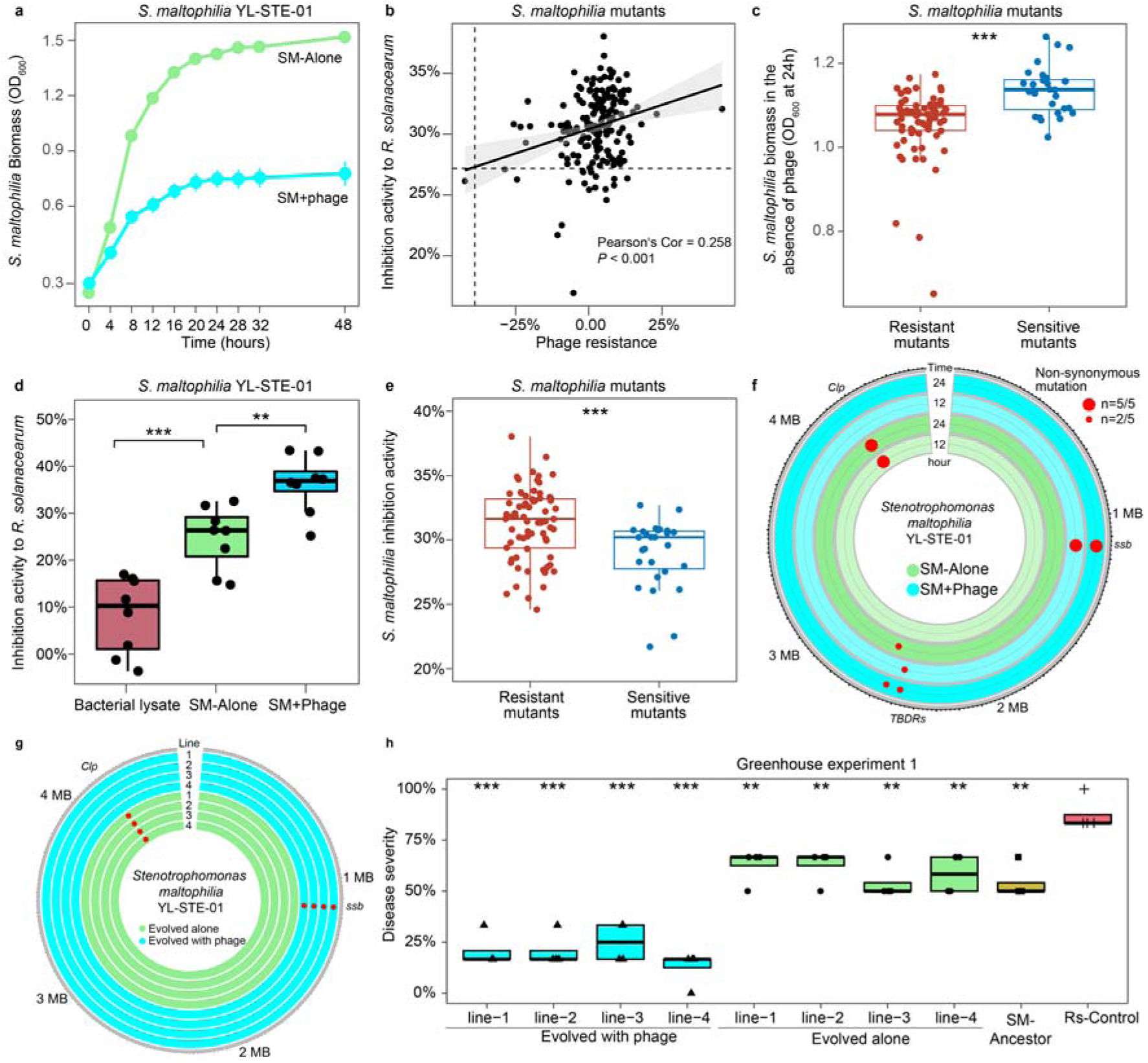
Co-evolution with phage increases both *S. maltophilia* phage resistance and *R. solanacearum* growth suppression. **a,** The growth curve of *S. maltophilia* YL-STE-01 in the presence (blue) and absence (green) of phage during laboratory selection experiment. Error bar shows standard deviation, *n = 8*. **b**, Correlation between the phage resistance and inhibition activity to *R. solanacearum* by each isolated mutant (mean, *n* = 3). The significance of linear regression is based on Pearson’s correlation coefficient. Black dashed lines indicate the value of ancestral *S. maltophilia*. **c,** Comparison of the growth (biomass) of ‘phage resistant mutants’ (*n* = 72) and ‘phage sensitive mutants’ (*n* = 28) at 24 h cultured in the absence of phage. Unpaired Wilcoxon test, \*\*\**P* < 0.001. **d**, Inhibition activity to *R. solanacearum* of *S. maltophilia* lysate and supernatant cultured in the absence (SM-Alone) or presence (SM+Phage) of phage (Unpaired t-test, \*\*\**P* < 0.001, \*\**P* < 0.01, *n* = 8). **e,** Comparison of the inhibition activity of *R. solanacearum* by phage resistant (*n* = 72) and phage sensitive (*n* = 28) *S. maltophilia* mutants at 24 h. Unpaired Wilcoxon test, \*\*\**P* < 0.001. **f**, Parallel mutations observed in laboratory evolved *S. maltophilia* populations in the presence (light blue rings) or absence (light green rings) of phages at 12 and 24 hours (*n* = 5). Points show parallel non-synonymous mutations that occur two or more times across treatment replicates in whole population sequencing of communities in given loci. Point sizes show the number of independent replicate populations in which the mutations were observed within treatment across a total of five biological replicates. The mutations observed at both 12 and 24 hours are labeled with the annotated gene functions. **g**, Mutations of greenhouse evolved *S. maltophilia* mutants identified across different loci in the presence (light blue rings) or absence (light green rings) of phages. Points show parallel non-synonymous mutations in given loci. **h,** Disease severity of tomato plants infected with *R. solanacearum* treated with water (Rs-Control), *S. maltophilia* ancestor (SM-Ancestor) and four *S. maltophilia* mutants evolved in the presence (Evolved with phage) and absence (Evolved alone) of *S. maltophilia*-specific phage (one mutant per replicate; four replicates from both treatments tested; Unpaired t-test compared with ‘Rs-Control’; *R. solanacearum* alone).

To explore if phage resistance could change the secretion of potential antimicrobial compounds by *S. maltophilia*, we extracted the supernatants from *S. maltophilia*-only and *S. maltophilia*-phage treatment replicates after 48h of selection and directly tested the suppressiveness of supernatants for *R. solanacearum* growth. Even though the presence of phage reduced *S. maltophilia* densities during the selection experiment, the suppressiveness of *S. maltophilia*-phage populations’ supernatant increased by 46.18% compared to *S. maltophilia*-only control populations’ supernatant (Unpaired t-test, *P* = 0.003, Fig. 1d). To eliminate the inhibitory effect of potential intracellular metabolites released from *S. maltophilia* cells due to phage lysis, we tested the suppressiveness of ancestral *S. maltophilia* cell lysate on *R. solanacearum* growth. We found that the cell lysate had much weaker effect compared to supernatants extracted from the *S. maltophilia*-only control and *S. maltophilia*-phage treatments (Unpaired t-tests, *P* < 0.001, Fig. 1d), suggesting that the phage-induced suppressiveness was likely due to compounds secreted by *S. maltophilia*.

To determine this inhibitory effect at the strain level, we tested the suppressiveness of *S. maltophilia* colonies isolated from phage treatment replicates on *R. solanacearum* growth. A total of 67.39% (124/184) of colonies showed higher *R. solanacearum* suppression compared to ancestral *S. maltophilia* strain, while only 2.17% (4/184) of colonies showed reduced suppression (Unpaired t-test, *P* < 0.05, Fig. 1b, Supplementary data S1; the remaining 56/184 colonies did not show significant change). Overall, phage resistance correlated positively with *R. solanacearum* growth suppression (lineal model, *P* < 0.001, Fig. 1e), and 65.22% of colonies (120/184) showed both increased phage resistance and pathogen suppressiveness (Unpaired t-test, *P* < 0.05, Fig. 1e, Supplementary data S1). We also found that phage-resistant colonies showed significantly higher pathogen suppression compared to phage-sensitive colonies (Unpaired Wilcoxon test, *P* < 0.001, Fig. 1e). In contrast, colonies isolated from the *S. maltophilia*-only treatment showed similar level of phage resistance (average of +1.65%) and pathogen suppression (average of +0.36%) as the ancestral *S. maltophilia* strain (Supplementary Table S1). Together, these results suggest that *S. maltophilia* could rapidly evolve resistance to its phage and that the evolution of phage resistance consistently led to increased *R. solanacearum* suppression, indicative of positive correlation between phage resistance and antagonism.

### Genetic basis of *S. maltophilia* phage resistance and disease suppressiveness

To explore the genetic and molecular basis of increased *S. maltophilia* phage resistance and suppressiveness to *R. solanacearum*, we used population sequencing to identify *S. maltophilia* variants enriched by phage selection during the selection experiment. Only a few SNPs were observed in the presence of phage at both 12h and 24h of sampling time points. We observed parallel mutations in the *Clp* gene encoding CRP-like protein (5/5 replicates, fixation index = 0.5, Fig. 1f, Supplementary Fig. S2) in the *S. maltophilia*-only control treatment populations. The *Clp* gene has previously been linked with temperature adaptation in *S. maltophilia* via c-d-GMP signalling^18^, but it is unclear what role it played in our experimental conditions.

In contrast, some non-synonymous mutations in *ssb* (5/5 replicates, fixation index = 0.781) and TonB-dependent receptor (*TBDR)* genes (2/5 replicates, fixation index = 0.125) were only observed in the *S. maltophilia*-phage treatment (Fig. 1f, Supplementary Fig. S2). The *ssb* gene encodes single-stranded DNA-binding protein and has previously been reported to be crucial in maintaining genome stability and stimulation of genetic recombination^19^. Although bacterial *ssb* protein can interact with numerous proteins involved in DNA metabolism^19^, it has not been directly linked to phage resistance or secretion of antimicrobial compounds previously. Instead, *TBDRs* gene has previously been reported as a *S. maltophilia* phage receptor-related gene^20^, suggesting it could be linked with phage resistance in *S. maltophilia*. However, it remains unclear how *TBDR* gene could have altered *S. maltophilia* disease suppressiveness.

To test if phage-mediated changes in *S. maltophilia* antimicrobial activity could enhance the suppression of *R. solanacearum*-mediated bacterial wilt disease *in planta*, a subset of colonies from *S. maltophilia*-only and *S. maltophilia*-phage treatments along with *S. maltophilia* ancestral strain were selected for tomato greenhouse experiment (*n* = 8, Supplementary Table S1; both the phage resistance and inhibition activity to *R. solanacearum* were clearly higher with isolates originating from *S. maltophilia*-phage compared to *S. maltophilia*-only control treatment). We found that phage resistant *S. maltophilia* isolates reduced the bacterial wilt disease severity of tomato by an average of 66.67% compared to phage-sensitive isolates from *S. maltophilia*-only control treatment (Unpaired t-test, *P* < 0.001, Fig. 1h). Moreover, the disease suppression of ancestral strain and isolates originating from *S. maltophilia*-only control treatment did not differ from each other (an average reduction of 35.12%; unpaired t-test, *P* > 0.05, Fig. 1h). To understand the underlying genetics, we re-sequenced the eight selected mutants using ancestral *S. maltophilia* strain as reference. In accordance with population sequencing results, all colonies from *S. maltophilia*-only treatment showed nonsynonymous mutations in the *Clp* gene, while all colonies from *S. maltophilia*-phage treatment showed nonsynonymous mutations in the *ssb* gene (Fig. 1g). None of the phage treatment colonies had mutations in the *TBDRs* gene, which aligns well with the population sequencing results where *ssb* mutations were found more frequently than *TBDR* mutations. Together, these results suggests that phage selection led to clear genomic divergence in *S. maltophilia* populations and that these changes took place only in a few loci. Moreover, while *Clp* mutation was likely linked to media adaptation, changes in phage resistance and pathogen suppression were likely caused by mutations in *ssb* and *TBDRs* genes.

### Phage selection alters the *S. maltophilia* transcriptional activity associated with anti-phage defense systems and secondary metabolism

To better understand how phage selection resulted in increased phage resistance and pathogen suppression, we compared the transcriptional responses of *S. maltophilia* in the presence and absence of phage at the 12h and 24h time points of the laboratory selection experiment. A total of 118 and 49 genes were up and down regulated, respectively, in the presence of phage relative to *S. maltophilia*-only control treatment at 12h sampling time point (Supplementary Fig. S3). Qualitatively similar transcriptomic responses were observed at 24h time point, but the number of differentially expressed genes was overall higher (182 up-regulated and 642 down-regulated genes in the presence vs. absence of phage; Fig. 2a). Specifically, we found that phage selection led to down-regulation of pathways associated with flagellar assembly, bacterial chemotaxis, biofilm formation and two-component signal transduction system, while fructose and mannose metabolism, 2-Oxocarboxylic acid metabolism, valine, leucine and isoleucine biosynthesis and type IV secretion system were significantly up-regulated (GSEA, *FDR* < 0.05, Fig. 2b).

**Figure 2.**
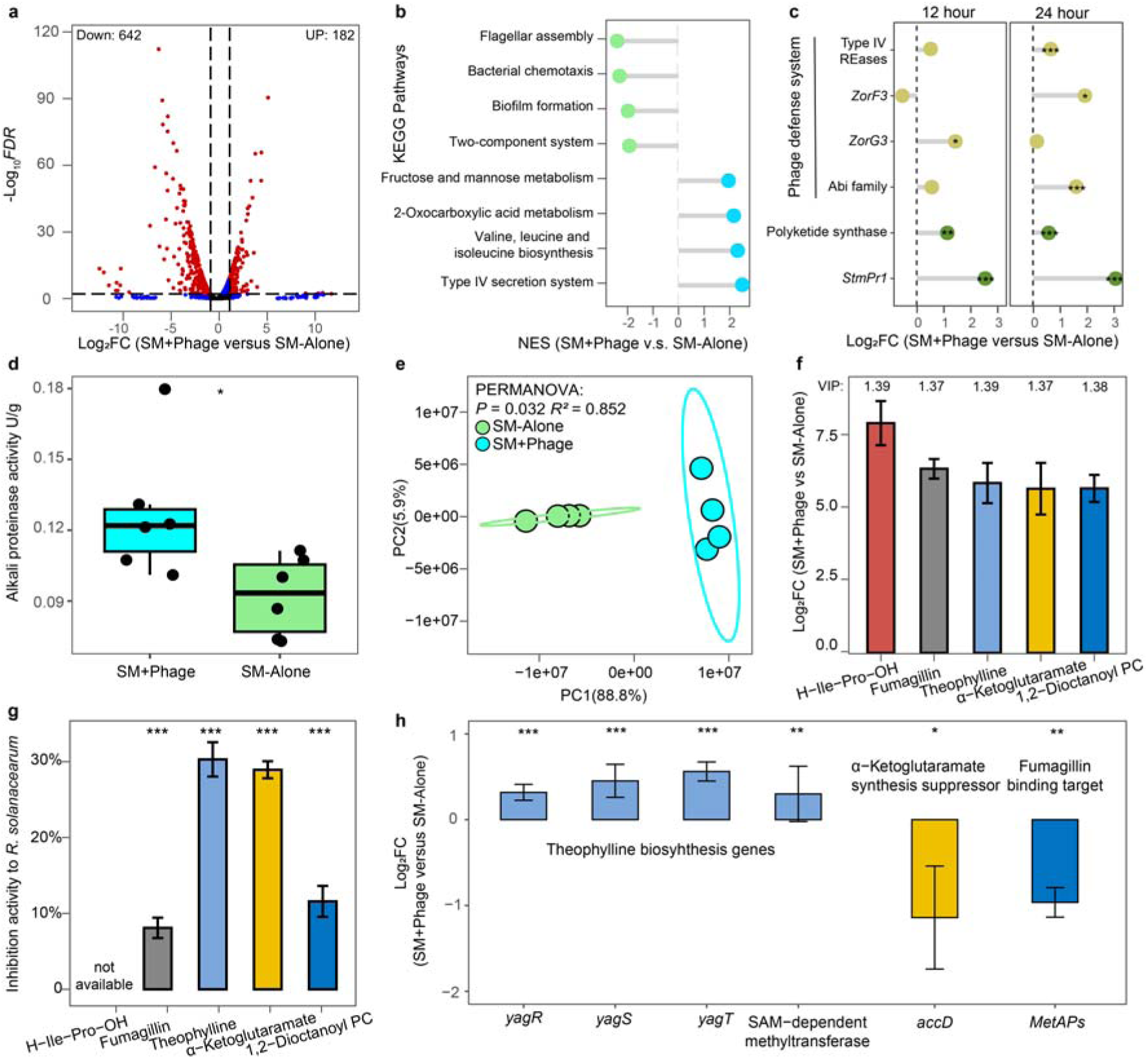
The changes in *S. maltophilia* gene expression and production of secondary metabolites due to phage selection during laboratory evolution experiment. **a,** Differentially expressed *S. maltophilia* genes in the presence of phage at 24-hour sampling time point. Red points denote significantly up or down regulated genes with *FDR* < 0.01 and |Log_2_ Fold Change| >1 (DEseq, *n* = 5). **b**, Normalized enrichment score (NES) of differentially expressed pathways in the presence of phage clustered by KEGG database at 24-hour sampling time point (GSEA, *FDR* < 0.05, *n* = 5). **c**, Fold change of genes related to phage defence systems (Type IV REases, *ZorF3*, *ZorG3* and Abi family) and antibiosis biosynthesis genes (Polyketide synthase and *StmPr1*) in the presence of phage (DEseq, ***: *FDR* < 0.001, **: *FDR* < 0.01, *: *FDR* < 0.05, *n* = 5). **d,** The extracellular alkali proteinase activity of *S. maltophilia* after 48-hour culturing in the presence (SM+Phage) and absence (SM-Alone) of *S. maltophilia*-specific phage. *: *P* < 0.05, unpaired t-test, *n* = 6. **e**, Comparison of *S. maltophilia* metabolome composition in the presence (light blue) and absence (light green) of phage at 24 hours (PCA, *n* = 4,). **f**, Fold change (mean ± SD) of top five up-regulated compounds in *S. maltophilia* supernatant in the presence of phage (Log transformed). The variable important projection (VIP) scores are labeled above each metabolite. g, *R. solanacearum* inhibition (mean ± SD) by the top five up-regulated chemical standards (Unpaired t-test compared to mono culture control, \*\*\**P* < 0.001, n.s.: *P* > 0.05, *n* = 6). **h,** Bar plot shows the mean (±SD) fold change of gene expressing data between ‘SM+Phage’ and ‘SM-Alone’ from RNA-seq at 24-hour. Each bar represents a gene related to the biosynthesis or suppression of theophylline (*yagR*, *yagS*, *yagT*, SAM-dependent methyltransferase), α-ketoglutaric acid (*accD*) and fumagillin (*MetAPs*). Statistical significances were determined by DEseq, ***: *FDR* < 0.001, **: *FDR* < 0.01, *: *FDR* < 0.05, *n* = 5.

In addition to mutations in potential phage receptor genes, phage resistance could be mediated by up-regulation of anti-phage defense systems. To explore this, we identified a total of 18 genes related to phage defense systems in *S. maltophilia* genome using both DefenseFinder^21^ and PADLOC^22^. Only *ZorG3* (part of Zorya defense system) was up-regulated in the presence of phage at 12-hour sampling time point while Type_IV_REases (part of Restriction-Modification system), GajA (part of Gabija system), ZorF3 (part of Zorya defense system) and Abi_2 (part of abortive infection system) were up-regulated in the presence of phage at 24-hour sampling time point (DEseq, *FDR* < 0.05, Fig. 2c, Supplementary Fig. S4a, Supplementary data S2). Together, these results indicate that phage presence upregulated multiple phage defense systems and metabolism-associated genes in *S. maltophilia* which could be associated with both phage resistance and production of antimicrobial compounds.

To study changes in pathogen suppression further, we analyzed how phage selection changed the expression of genes associated with secondary metabolism (genes belonging to ‘secondary metabolite biosynthesis, transport and catabolism’; category Q in COG database), which could directly explain the observed increase in *R. solanacearum* suppression. We found that only six out of 52 secondary metabolism-related genes were differentially expressed in the presence of phage at 12-hour sampling time point (DEseq, *P* < 0.05, Supplementary Fig. S4b, Supplementary data S2) with half of these genes showing up-regulation (*DsbA_FrnE*, *Tam* and *MlaE*). At 24h sampling time point, 20 secondary metabolism-related genes were differentially expressed in the presence of phage (DEseq, *P* < 0.05, Supplementary Fig. S4b, Supplementary data S2) with seven genes showing up-regulation. Specifically, the *DsbA_FrnE* polyketide synthase gene was significantly upregulated in the presence of phage at both time points (DEseq, *FDR* < 0.05, Fig. 2c, Supplementary data S2), which has previously been linked to bacterial wilt disease suppression in soils^23^. We also found that an extracellular protease *StmPr1* gene, which encodes the alkaline serine proteolytic enzyme^24^, was clearly upregulated in the presence of phage at both time points (DEseq, *FDR* < 0.001, Fig. 2c, Supplementary data S2). As previous studies have shown that alkaline serine proteolytic enzyme of *S. maltophilia* can directly inhibit *R. solanacearum*^25^, we analyzed the *S. maltophilia* supernatant extracted at 48h time point of the selection experiment and found that the phage selection significantly increased the alkali proteinase activity relative to *S. maltophilia*-only control treatment (Unpaired t-test, *P* = 0.029, Supplementary Fig. 2d).

Interestingly, although *ssb* gene was mutated under phage infection, its transcriptional expression was not significantly changed. As *ssb* protein can bind to other proteins, we combined PDBePISA^26^ and PRODIGY^27^ to infer the protein-protein interactions between ancestral and mutated *ssb* proteins with the up-regulated phage defense and secondary metabolite proteins using structure predictions based on Boltz-2^28^. Results showed the phage-driven mutations in *ssb* gene leads to overall larger interface area, lower binding affinity energy and more intermolecular contacts when binding with proteins of Abi family, Type IV REases, *StmPr1* and polyketide synthase (Supplementary Table S2). This indicates mutated *ssb* protein had potentially higher potential to interact with identified proteins related to phage resistance and secondary metabolism. Together, these results suggest that phage selection upregulated the expression of genes associated with secondary metabolism in *S. maltophilia*, explaining the observed increase in *R. solanacearum* growth inhibition.

To link changes in *S. maltophilia* gene expression with changes in secreted metabolites, we conducted metabolomic analysis at 24h sampling time point. The phage selection significantly changed the *S. maltophilia* metabolome composition (PERMANOVA, *P* = 0.032, *R*^2^ = 0.852, Fig. 2e) and 228 out of 1286 identified compounds were produced at significantly higher levels in the presence of phage (Unpaired t-test, Log_2_FoldChange > 1, *P* < 0.05). The top five compounds that were produced relatively more in the presence of phage included H-Ile-Pro-OH (255.43-fold), fumagillin (82.85-fold), theophylline (60.42-fold), α-ketoglutaric acid (53.91-fold), and 1,2-Dioctanoyl PC (52.22-fold) (Fig. 2f). To validate if these compounds were linked with pathogen suppression, we tested four out of five compounds for their antimicrobial activity against *R. solanacearum* using available chemical standards (H−Ile−Pro−OH was not available and could not be validated). All tested compounds significantly inhibited the *R. solanacearum* growth and theophylline and α-ketoglutaric acid showed the highest antimicrobial activity (Unpaired t-test, *P* < 0.05, Supplementary Fig. 2g). To further link metabolomic variation with transcriptomics data, we compared the expression of genes involved in the biosynthesis and catabolism pathways of subset of these compounds. The theophylline is normally methylated from xanthines^29^ and we found that three genes encoding xanthine dehydrogenase (*yagR*, *yagS*, *yagT*) as well as one gene encoding SAM-dependent methyltransferase were significantly over-expressed under phage selection at 24 h sampling time point (DEseq, *FDR* < 0.05, Fig. 2h). We also found that acetyl-CoA carboxylase carboxyl transferase subunit beta (*accD*) gene, which suppresses α-ketoglutaric acid synthesis^30^, and methionine aminopeptidase gene (*MetAPs*), which has been identified as the binding target of fumagillin^31^, were both significantly down-regulated in the phage treatment at 24 h sampling time point (DEseq, *FDR* < 0.05, Fig. 2h, Supplementary data S2). These findings suggest that down regulation of these genes could have led to increased accumulation of theophylline, α-ketoglutaric acid and fumagillin in the supernatant in the presence of phage. Together, these results demonstrate that selection by phage can rewire the *S. maltophilia* gene expression and secondary metabolism, leading to increased production of antimicrobial compounds that can suppress *R. solanacearum* growth as a by-product.

### *S. maltophilia* can evolve phage resistance via *ssb* mutations in the plant rhizosphere

To test if the ancestral *S. maltophilia* can also evolve phage resistance in the tomato rhizosphere, we conducted another greenhouse experiment. To achieve this, we inoculated *S. maltophilia* YL-STE-01 ancestral bacterial strain either alone or together with its phage YL-STE-01P into sterilized soil with tomato seedlings in the absence of *R. solanacearum* (four replicate plants per treatment). After 10-days, we collected the rhizosphere soil and isolated one representative *S. maltophilia* colony from each plant replicate from both treatments (4 colonies per treatment, Fig. 3a). We found that colonies isolated from *S. maltophilia*-phage treatment had evolved both higher *R. solanacearum* growth suppression (46.48% relative increase, *P* = 0.008) and phage resistance (94.08% relative increase, *P* < 0.001) compared to colonies isolated from *S. maltophilia*-only treatment (Unpaired t-test, Fig. 3b-c).

**Figure 3.**
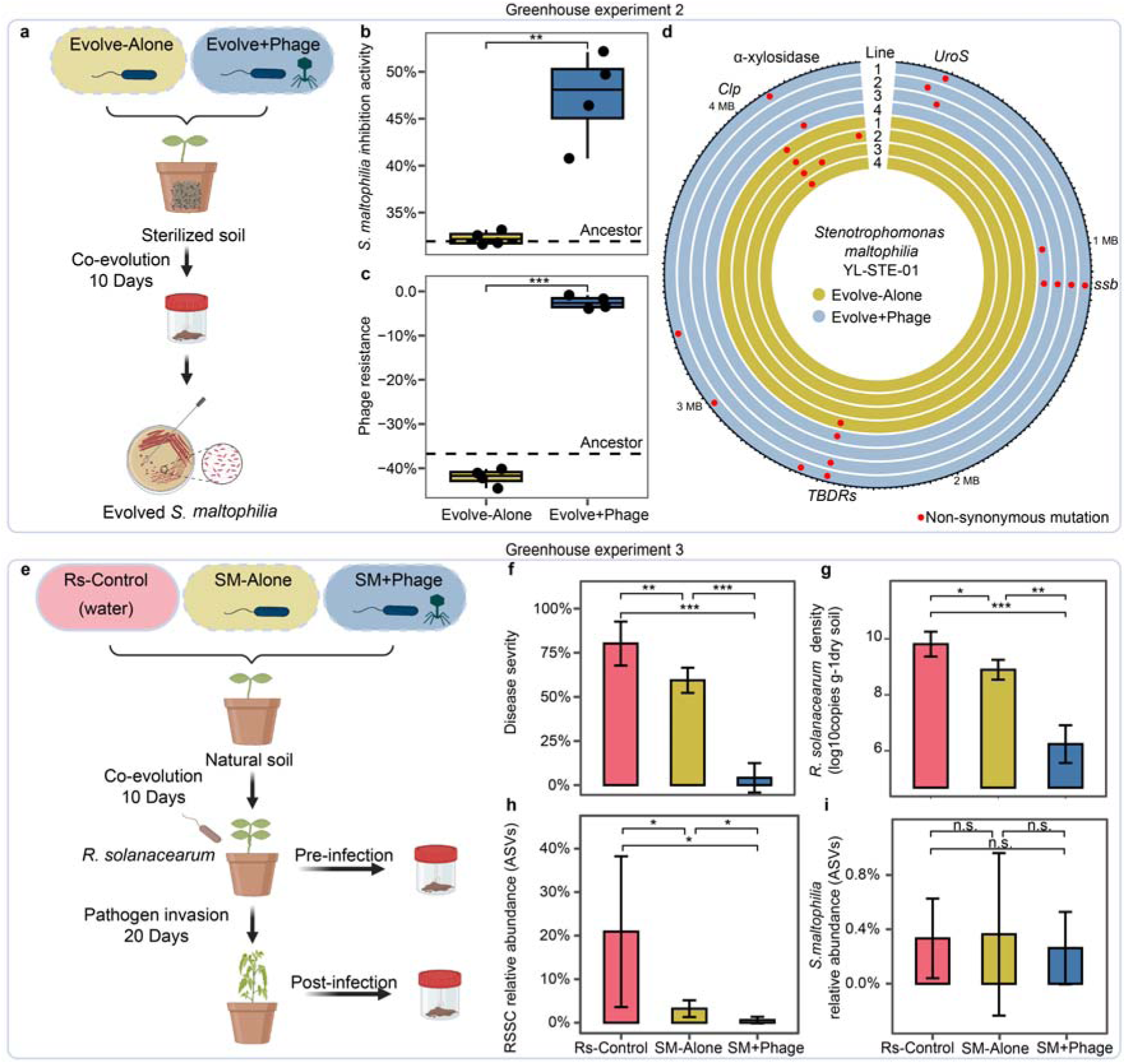
Phage-*S. maltophilia* co-culturing in the rhizosphere selects for phage resistant mutants and enhances microbiome disease suppressiveness. **a,** Schematic diagram for greenhouse experimental design to determine *S. maltophilia* phage resistance evolution in the rhizosphere by isolating *S. maltophilia* strains from the tomato rhizosphere treatment without or with *S. maltophilia* phage. b-c, The *R. solanacearum* inhibition activity (b) and phage resistance (c) of *S. maltophilia* colonies that evolved in the absence (Evolve-Alone) or in the presence of phage (Evolve+Phage) during the greenhouse experiment (Unpaired t-test, \*\*\**P* < 0.001, \*\**P* < 0.01, *n* = 4). **d**, Mutations identified across different loci of *S. maltophilia* mutants that evolved in the absence (Evolve-Alone) or in the presence of phage (Evolve+Phage) during the greenhouse experiment (*n* = 8). Points show parallel non-synonymous mutations in given loci. The mutations observed in at least two colonies are labeled with the annotated gene functions. **e**, Schematic diagram for greenhouse experimental design to assess the effect of *S. maltophilia*-phage treatment on disease progressions and *R. solanacearum* and *S. maltophilia* densities. The microbes of *S. maltophilia* alone (SM-Alone), *S. maltophilia* with its phage (SM+Phage) and water (Rs-Control) treatments were inoculated to tomato rhizosphere simultaneously at the beginning of the experiment. After 10 days, rhizosphere samples were collected at ‘pre-infection stage’. *R. solanacearum* pathogen was then inoculated in all treatment and after 20 days, rhizosphere samples were collected at ‘post-infection stage’. f-i, Disease severity (mean ± SD, f), *R. solanacearum* density (mean ± SD, based on qPCR of the *fliC* gene, g), *R. solanacearum* (h) and *S. maltophilia* relative abundance (mean ± SD, based on amplicon sequencing data, i) of tomato plants treated with *S. maltophilia* alone (SM-Alone), *S. maltophilia* with its phage (SM+Phage) and water (Rs-Control) at the ‘post-infection stage’. Unpaired t-test, n.s.: *P* > 0.05, \**P* < 0.05, \*\**P* < 0.01, \*\*\**P* < 0.001, *n* = 4. Panels a and e were created with BioRender.com.

We also re-sequenced the colonies to identify mutations relative to ancestral *S. maltophilia* strain. In line with previous lab experiments, all colonies from *S. maltophilia*-only treatment showed nonsynonymous mutations in the *Clp* gene, while all colonies from *S. maltophilia*-phage treatment showed nonsynonymous mutations in the *ssb* gene. In addition, three out of four colonies in *S. maltophilia*-phage treatment had nonsynonymous mutations in *TBDRs* and a few additional mutations, which were not observed during laboratory experimental evolution (Supplementary Data S3). For example, two colonies from *S. maltophilia*-phage treatment showed nonsynonymous mutations in the uroporphyrinogen-III synthase (*UroS)* gene and one colony from *S. maltophilia*-phage and *S. maltophilia*-only treatments showed nonsynonymous mutations in the α-xylosidase gene (Fig. 3d). As α-xylosidase is involved in xyloglucan degradation, this mutation could have been adaptive in the rhizosphere. This experiment demonstrate that *S. maltophilia* can rapidly evolve increased phage resistance and pathogen suppression in the tomato plant rhizosphere likely through mutations in the *ssb* and *TBDR* genes.

### *S. maltophilia*-specific phage enhances the suppression of *R. solanacearum* in natural soil

To understand the effects of *S. maltophilia*-specific phage on the *R. solanacearum* growth suppression in the presence of rhizosphere microbiota, we conducted a separate greenhouse experiment with tomato in natural soil (Fig. 3e) inoculated with ancestral *S. maltophilia* strain without and with its phage. The *S. maltophilia* strain and its phage were inoculated into the rhizosphere at the tomato seeding stage (sterilized water with no inoculated microbes was used as negative control), and after 10 days, we collected rhizosphere soil samples from all treatments and designated them as ‘pre-infection stage’ samples. All treatment replicates were then inoculated with phytopathogenic *R. solanacearum* (Fig. 3e; see methods) and sampled again after 20 days (‘post-infection stage’ samples). We found that the levels of disease incidence had stabilized by sampling day 20, and that the highest levels of disease were observed in *R. solanacearum*-only treatment, resulting in an average of 79.16% disease severity (Fig. 3f). In line with our previous study^3^, inoculation of *S. maltophilia* clearly reduced the disease symptoms by 15.79% (Unpaired t-test, *P* = 0.035, Fig. 3f) relative to the *R. solanacearum*-only control treatment. Strikingly, the co-inoculation of *S. maltophilia* and its phage decreased the disease severity most clearly (94.74% reduction relative to the *R. solanacearum*-only control treatment; Unpaired t-test, *P* < 0.001, Fig. 3f), indicative of prophylactic treatment effect. The ‘post-infection stage’ samples were also used to compare *R. solanacearum* cell densities between treatments using qPCR. In line with reduced disease severity, inoculating plants with *S. maltophilia* reduced pathogen densities by 9.34% (Unpaired t-test, *P* = 0.01, Fig. 3g), while the co-inoculation of *S. maltophilia* and its phage decreased pathogen density more drastically by 36.41% (Unpaired t-test, *P* < 0.001, Fig. 3g) relative to the *Ralstonia*-only control treatment. We also analyzed changes in *R. solanacearum* and *S. maltophilia* abundances from ‘post-infection stage’ samples using amplicon sequence data (relative 16S rRNA copy numbers). The total abundances of ASVs belonging to *Ralstonia solanacearum* species complex (RSSC, a combination of ASV2, ASV4, ASV44 and ASV8022, Supplementary Fig. S5a) were reduced by 84.58% in *S. maltophilia*-only and by 97.08% in *S. maltophilia*-phage treatments compared to *R. solanacearum*-only treatment (Unpaired t-test, *P* < 0.001, Fig. 3h). Interestingly, the presence of *S. maltophilia*-specific phage did not cause significant reduction in *S. maltophilia* ASV abundances (a combination of ASV3, ASV15 and ASV18, Supplementary Fig. S5b). This suggests that phage did not trigger strong density control of *S. maltophilia* (Unpaired t-test, *P* > 0.05, Fig. 3i), potentially because the length of the experiment allowed *S. maltophilia* phage resistance evolution as observed in earlier greenhouse experiments. Together, these results suggest that *S. maltophilia* can protect plants from *R. solanacearum* infections in natural soil by reducing pathogen densities, and that this effect was stronger in the presence of *S. maltophilia* and its phage.

### *S. maltophilia*-specific phage stabilizes rhizosphere bacterial diversity, community composition and species co-occurrence networks

To determine the potential indirect effects of *S. maltophilia*-specific phage on rhizosphere microbiota, we tested how *S. maltophilia* and its phage affected the diversity and composition of rhizosphere microbiota in natural soil by comparing ‘post-infection stage’ samples with ‘pre-infection stage’ samples (Fig. 3e). We first identified bacteria that were enriched in each treatment. A total of six out of 42 identified bacterial genera were specifically enriched in *R. solanacearum*-only treatment. These genera included *Ralstonia*, which reached 5.45% relative abundance in the bacterial community (Turkey’s HSD test, Fig. 4a, Supplementary Data S4). Interestingly, the co-inoculation of *R. solanacearum* and *S. maltophilia* did not clearly change the relative abundances of any bacterial genera. However, the co-inoculation of *R. solanacearum* with *S. maltophilia* and its specific phage led to enrichment of nine bacterial genera, which included several previously reported disease-suppressive taxa such as *Massilia*^32^, *Micromonospora*^33^, *Nocardioides*^5^ and *Sphingomonas*^34^ (Fig. 4a, Supplementary Data S4). Specifically, the presence of *S. maltophilia*-specific phage changed the abundances of 341 ASVs, of which 63.05% (*n* = 215) were enriched (Unpaired t-test, *P* < 0.05, Fig. 4b, Supplementary Data S5). While the microbiome α-diversity (Shannon index) decreased by 10.5% in *R. solanacearum*-only treatment, no reduction in α-diversity was observed in *R. solanacearum* and *S. maltophilia* co-inoculation treatment (100.19%). However, including *S. maltophilia*-specific phage to *R. solanacearum* and *S. maltophilia* co-inoculation treatment significantly increased the microbiome α-diversity by 11.24% (Unpaired t-test, *P* = 0.019, Fig. 4c). When comparing β-diversity, the largest shift in microbiome composition was observed in *R. solanacearum*-only treatment compared to the other treatments (PERMANOVA and unpaired t-test, *P* < 0.05, Fig. 4d-e, Supplementary Table S3). Moreover, the community composition of *R. solanacearum*-only treatment differed significantly from both *S. maltophilia* treatments (PERMANOVA, *P* < 0.05, Fig. 4d-e, Supplementary Table S3), while the presence of *S. maltophilia*-specific phage maintained the microbial community composition close to “pre-infection stage” sample levels (PERMANOVA, *P* > 0.05, Fig. 4d, Supplementary Table S3).

**Figure 4.**
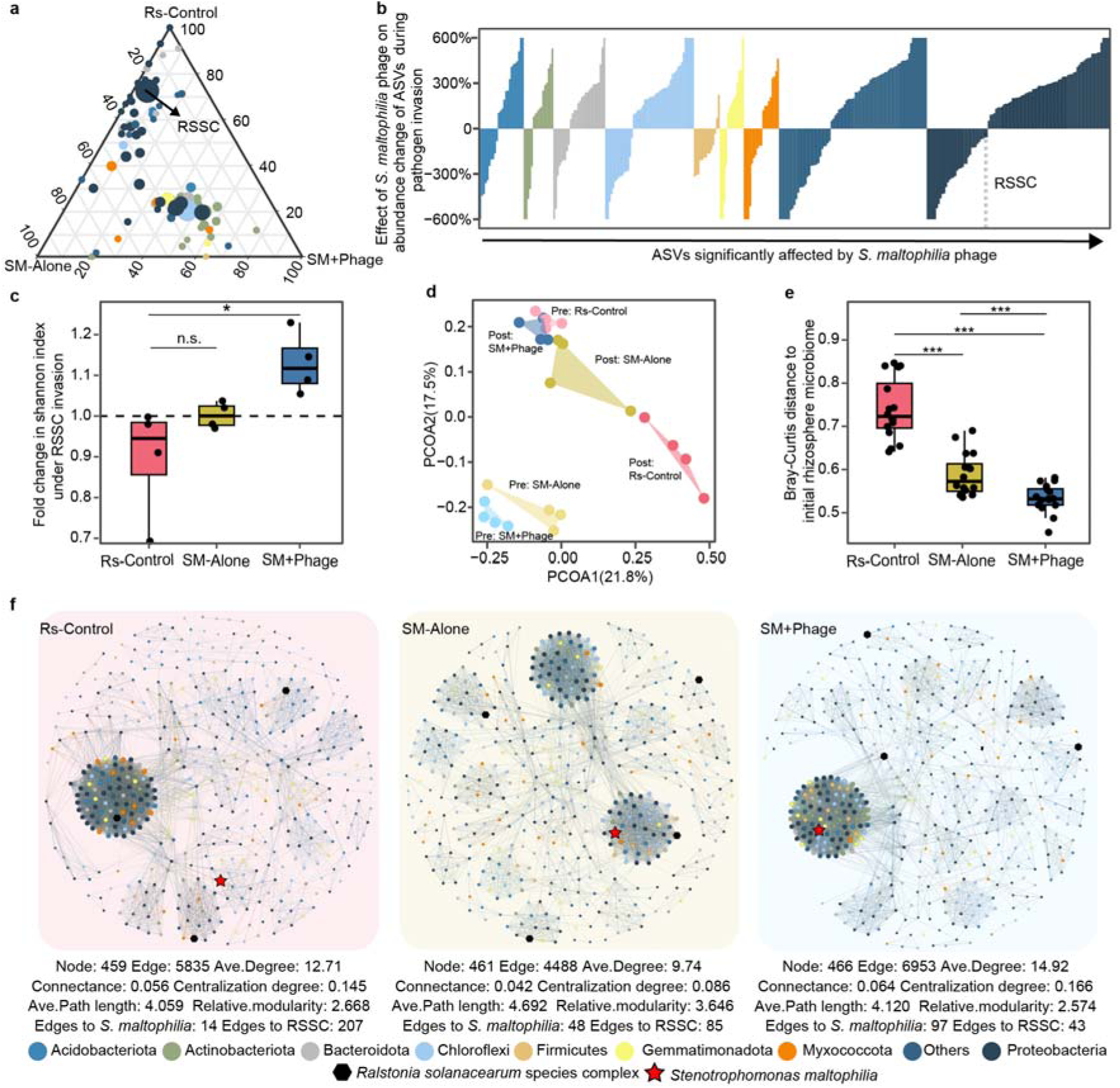
*S. maltophilia*-phage stabilizes microbiome diversity and composition under *R. solanacearum* invasion. **a**, Compositional differences of tomato rhizosphere microbiomes under *R. solanacearum* invasion when treated with *S. maltophilia* alone (SM-Alone), *S. maltophilia* with its phage (SM+Phage) or water (Rs-Control; *R. solanacearum* only) at ‘post-infection stage’. Only genera varying significantly among treatments are shown (Tukey’s honestly significant difference test after one-way ANOVA). **b**, Average abundance (ASV) changes during pathogen invasion due to presence of *S. maltophilia* phage. **c**, Fold change of microbiome Shannon index during *R. solanacearum* invasion between different treatments. Unpaired t-test, \**P* < 0.05, n.s.: *P* > 0.05, *n* = 4. **d**, PCOA showing changes in bacterial community structure between different treatments at ‘pre-infection stage’ (Pre) and ‘post-infection stage’ (Post). **e**, Bray-Curtis distance of microbiome community composition compared with the initial microbiome. Unpaired t-test, \*\*\**P* < 0.001, *n* = 16. **f**, Networks of major ASVs (relative abundance > 0.02% and present in at least half of samples) based on pairwise maximal information coefficient (mic > 0.8). Each network was generated by combining samples from both pre-infection and post-infection stages (*n* = 8). The network properties are given below networks. ASVs belonging to RSSC and *S. maltophilia* are highlighted with black hexagons and red stars, respectively.

To further analyze treatment differences in bacterial community composition, ASV-level co-occurrence networks were constructed for both sampling time points and pairwise correlations between ASVs between treatments were compared based on the maximal information coefficient (MIC). We found that ASVs in *R. solanacearum-S. maltophilia*-phage treatment formed the most complex network with the highest edge number (6953), average degree (14.92), connectance (0.064) and centralization degree (0.166) values, compared to other two pathogen treatments (Fig. 4f). The *R. solanacearum*-only network contained one major module with *R. solanacearum*. In contrast, *R. solanacearum-S. maltophilia*-phage network formed two medium-sized modules with one containing both *S. maltophilia* and *R. solanacearum*, and the other module including *S. maltophilia* but not *R. solanacearum* (Fig. 4f). We further compared the sub-networks connected to *S. maltophilia* and *R. solanacearum* and found that the number of edges between *S. maltophilia* and other bacteria was higher in the presence (*n* = 97) versus absence (*n* = 48) of *S. maltophilia*-specific phage, while the number of edges between *R. solanacearum* and other bacteria was lower in the presence (n = 43) versus absence (n = 85) of phages (Supplementary Fig. S6). Together, these results highlight that *S. maltophilia*-specific phage can drive community-wide changes in rhizosphere microbiomes, leading to stabilization of microbiome community diversity, composition and species co-occurrence, which are often altered under *R. solanacearum* invasion^3,35^.

### *S. maltophilia*-specific phage improves the suppressiveness of other resident bacteria against *R. solanacearum*

To link changes in bacterial community composition with changes in microbiome-wide disease suppressiveness, we used the microbiome data to identify bacterial ASVs that had high potential to interact with *S. maltophilia* based on their 1) co-occurrence with *S. maltophilia* (*n* = 79, Criterion I) and 2) their enrichment in the presence of *S. maltophilia*-specific phage (*n* = 196, Criterion II, Fig. 5a). A total of 19 ASVs fulfilled these both criteria (Fig. 5a), and although most of these taxa were classified as species that are hard to culture, we could isolate two candidate bacterial species from the endpoint samples: *Enterobacter ludwigii* YL-ENT31 and *Pseudomonas baetica* PSE01, which were phylogenetically close to ASV432 (100%) and ASV192 (98.387%) in the microbiome dataset (based on 16S rRNA sequence, Supplementary Fig. S7a-b) and have previously been identified as a plant growth-promoting bacterial strains^3,36^. We first determined the metabolite-mediated interactions between *S. maltophilia* and these two bacterial species in controlled lab conditions. It was found that *E. ludwigii* supernatants inhibited the *S. maltophilia* growth (Unpaired t-test, *P* < 0.001, Supplementary Fig. S7c), while *S. maltophilia* supernatants facilitated the growth of *E. ludwigii* (Unpaired t-test, *P* < 0.001, Supplementary Fig. S7d), which suggests that *E. ludwigii* could exploit *S. maltophilia*. In contrast, *P. baetica* supernatants facilitated the growth of *S. maltophilia* (Unpaired t-test, *P* < 0.001, Supplementary Fig. S7e), while *S. maltophilia* supernatants inhibited the growth of *P. baetica* (Unpaired t-test, *P* < 0.001, Supplementary Fig. S7f), suggesting that *S. maltophilia* could exploit *P. baetica*. Interestingly, the presence of *S. maltophilia*-specific phage reduced both supernatant effects (Unpaired t-test, *P* < 0.001, Supplementary Fig. S7d,f), resulting in weaker antagonism between both bacterial species pair combinations, indicative of potential stabilizing effect that could potentially promote their co-existence.

**Figure 5.**
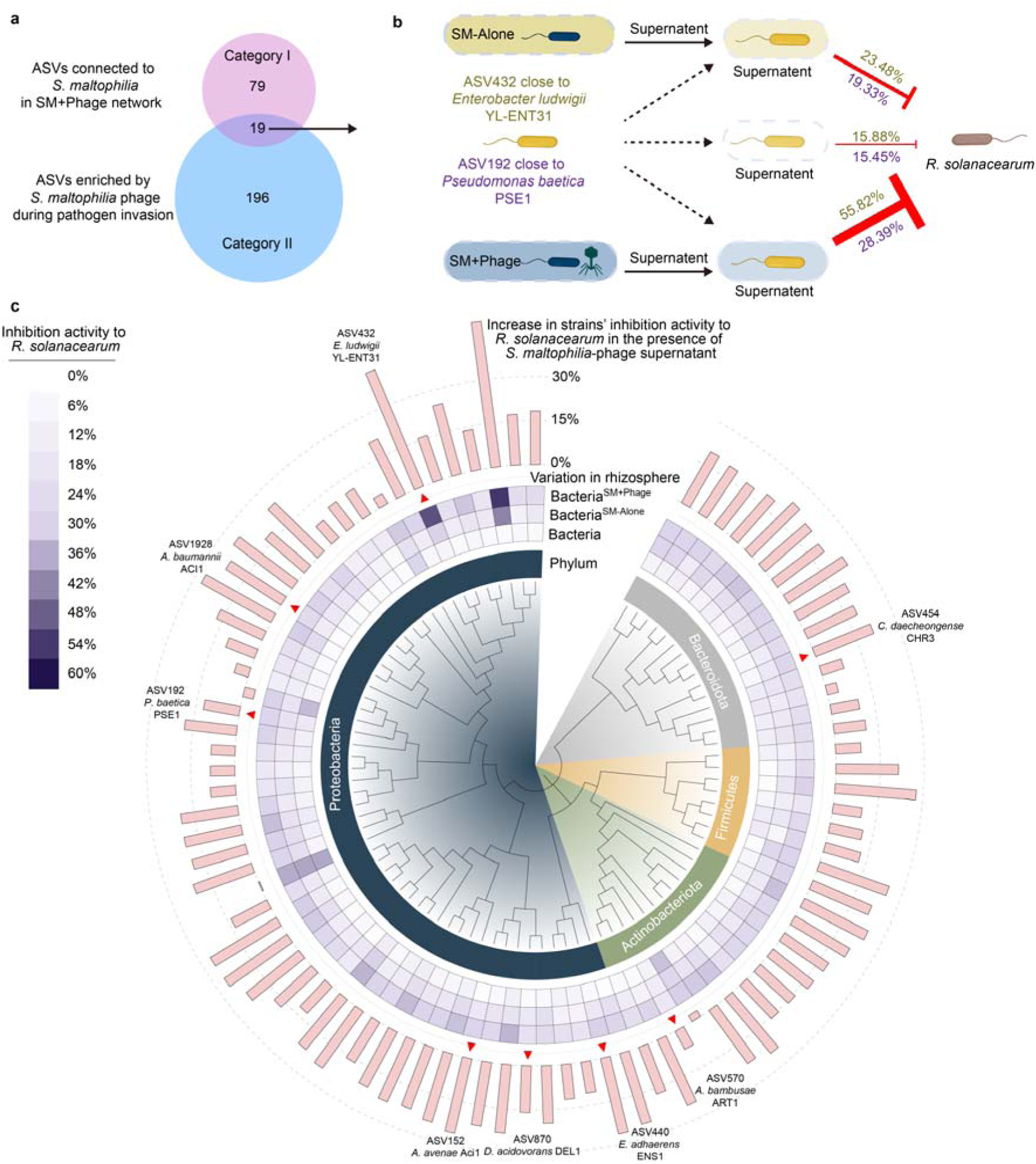
*S. maltophilia*-specific phage co-culture supernatants enhance the antagonism of resident bacteria to *R. solanacearum*. **a**, Venn diagrams displaying the overlap between ASVs connected to *S. maltophilia* in ‘SM+Phage’ network (Category I) and those enriched by *S. maltophilia* phage during pathogen invasion (Category II). **b**, Schematic diagram showing the effect of *S. maltophilia*-phage supernatants on the suppressiveness of *E. ludwigii* (dark red) and *P. baetica* (indigo) to *R. solanacearum*. Numbers indicate the inhibition activity based on the reduction of *R. solanacearum* biomass. See Fig.S9 for details. **c**, Effects of *S. maltophilia*-phage co-culture supernatant on *R. solanacearum* inhibition activity across different taxa. The phylogenetic neighbor-joining tree in the center shows the taxonomic relationships among 88 resident bacterial species classified belonging to four different phyla. The heatmap in the middle ring shows the inhibition activity of each bacterium to *R. solanacearum* treated with water (Bacteria) and *S. maltophilia* supernatant previously cultured in the absence (Bacteria^SM^) and presence (Bacteria^SM+Phage^) of phage (mean, *n* = 5). Eight bacteria are highlighted with triangles for their identity with ASVs whose relative abundance were increased in the presence of *S. maltophilia* phage during greenhouse experiment (Category II). The increase in strains’ inhibition activity to *R. solanacearum* in the presence of *S. maltophilia*-phage supernatant is shown as bars in the outer ring (mean, *n* = 5). The panel b was created with BioRender.com.

We next tested how interactions between the two resident rhizosphere bacteria, *S. maltophilia* and its phage affected the growth of *R. solanacearum* by using supernatant assays. We found that the inhibition of *R. solanacearum* was stronger by pairwise bacterial culture supernatants compared to *S. maltophilia* monoculture supernatant with both *E. ludwigii* (47.86%, Supplementary Fig. S7g) and *P. baetica* (25.11%, Supplementary Fig. S7h) (Unpaired t-test, *P* < 0.05, Fig. 5b). Crucially, this inhibitory effect was increased when the *S. maltophilia*-specific phage was included in the bacterial co-cultures, resulting in 251.51% (*E. ludwigii*, Supplementary Fig. S7g) and 83.75% (*P. baetica*, Supplementary Fig. S7h) increase in pathogen suppression (Unpaired t-test, *P* < 0.05, Fig. 5b). As the effects of *S. maltophilia* supernatants had much smaller inhibitory effects on *R. solanacearum* in the absence of *E. ludwigii* or *P. baetica*, the improved suppressiveness was likely caused by the metabolites produced by *E. ludwigii* and *P. baetica* (Supplementary Fig. S7g-h). Moreover, the *S. maltophilia*-specific phage could indirectly increase the suppressiveness of pairwise bacterial communities through interactions with *S. maltophilia*.

To understand the mechanisms behind the increased suppressiveness of the other resident bacteria, we focused on the *E. ludwigii* YL-ENT31. We first identified the *E. ludwigii* secondary metabolite biosynthesis gene clusters using antiSMASH (Supplementary Fig. S8a-b), identifying several siderophore biosynthesis clusters including Enterobactin and Aerobactin, which could also have antimicrobial potential^37^. We then compared the *E. ludwigii* metabolomes when cultured with *S. maltophilia*-only and *S. maltophilia*-phage supernatants. It was found that already *S. maltophilia*-only supernatant significantly changed the metabolome of *E. ludwigii* (*F* = 6.88) but that this effect became more pronounced in the presence of *S. maltophilia*-phage combination (*F* = 12.64, PERMANOVA, *P* < 0.05, Supplementary Fig. S8c). Among the identified compounds, ketoprofen acid, lactoferrin, dethiobiotin and traumatic acid were clearly enriched in the presence of phage (VIP-value > 1, *P* < 0.05, Supplementary Fig. S8d). Interestingly, lactoferrin is an iron-containing protein whose biosynthesis may possibility require the participation of bacterial siderophores^38^. Pure compounds of ketoprofen acid, lactoferrin, dethiobiotin and traumatic acid were further used to directly test their inhibitory activity on *R. solanacearum*. All tested compounds could significantly suppress the *R. solanacearum* growth with some of them having relatively larger effect than the others (ketoprofen acid: 34.71%; lactoferrin: 25.46%; dethiobiotin: 17.61% and traumatic acid: 4.34%; unpaired t-test, *P* < 0.05, Supplementary Fig. S8e). Together, these results demonstrate that especially when combined with its phage, *S. maltophilia* can indirectly increase microbiome-wide soil suppressiveness by activating production of antimicrobial compounds by other resident bacterial species.

Finally, we investigated the generality of our findings by experimentally testing if *S. maltophilia* and its phage could trigger increased suppression of *R. solanacearum* by 88 different resident rhizosphere bacterial species (covering four phyla) that were isolated from the same tomato field as the two representative resident bacterial strains (*E. ludwigii* and *P. baetica*; Supplementary data S6). We quantified the inhibition of *R. solanacearum* in the lab by using supernatants of each bacterium treated with water (negative control) or treated with *S. maltophilia* supernatants cultured in the absence and presence of phage instead of water. We found that 84.09% (74/88) of resident bacteria showed increased suppressiveness to *R. solanacearum* in the presence of *S. maltophilia*-phage supernatant compared to water control (Unpaired t-test, *P* < 0.05, Fig. 4c, Supplementary data S6). Moreover, the presence of *S. maltophilia*-specific phage improved the inhibition of 62.5% (55/88) by resident bacteria compared to when supernatants derived from *S. maltophilia*-only cultures were used (Unpaired t-test, *P* < 0.05, Fig. 5c, Supplementary data S6). Furthermore, we found that eight tested bacteria were phylogenetically similar to ASVs whose abundances were enriched by *S. maltophilia*-specific phage between ‘pre-infection stage’ and ‘post-infection stage’ samples during the greenhouse experiment (Fig. 5c, Supplementary data S7). Together, these results indicate that *S. maltophilia*-specific phage can indirectly trigger increased production of antimicrobial compounds by other rhizosphere bacteria, resulting in microbiome-wide soil suppressiveness against *R. solanacearum*.

## Discussion

Although phage therapy against pathogens have been developed for over a century^39^, the role of phages targeting non-pathogenic bacteria is still poorly understood in eukaryotic microbiomes. Here we explored the consequences of phage selection on probiotic *S. maltophilia* bacterium and discovered that evolution phage resistance correlates positively with increased suppression of phytopathogenic *R. solanacearum* bacterium via increased secretion of antimicrobial compounds. Furthermore, we found that *S. maltophilia*-specific phage could indirectly enrich other disease suppressing bacteria and increased their antimicrobial activity in the tomato rhizosphere, increasing the microbiome-wide disease suppressiveness. Together these findings suggest that phage resistance evolution does not always lead to trade-offs with other traits, and that single phage can have significant cascading effects on the rhizosphere microbiome diversity, composition and functioning in terms of enhanced disease suppression.

Increased phage resistance of *S. maltophilia* could be explained by potential resistance mutations and activation of several phage defense systems. Mutations in TonB-dependent receptors have previously been identified as *S. maltophilia* phage receptors and could have also have provided genetic phage resistance to *S. maltophilia* in our experiments where it was co-selected along with *ssb* mutations in the lab and in the rhizosphere. However, TonB mutations did not emerge consistently across all replicates in contrast to mutations in *ssb* gene, which were observed across all phage treatment replicates both in the lab and rhizosphere evolution experiment. The *ssb* gene encodes single-stranded DNA-binding protein and has not previously been linked with phage resistance. By using protein interaction predictions, we found that mutated version of *ssb* gene allowed broader and more efficient interactions with other proteins related to both phage defense systems and polyketide biosynthesis proteins, suggesting that it could have enhanced both phage resistance and biosynthesis of antimicrobials. While molecular and biochemical studies are still needed to directly link *ssb* gene mutations with observed phenotypic changes, our result suggest that *ssb* gene showed strong pleiotropic synergy with several other genes associated with phage resistance and antibiosis traits in *S. maltophilia*.

We also found that several phage defense systems were upregulated in the presence of phages simultaneously when the proportion of *ssb* mutant variants increased, including Zorya (ZorG3 and ZorF3), Resriction-Modification system (Type_IV_REases), Gabija (GajA) and abortive infection system (Abi_2). All these systems have been detected in *S. maltophilia* in a previous pangenomic study except for Zorya and Abi_2, which are part of AbiD, AbiD1 and AbiF abortive infection systems^40^. Of these systems, Zorya was upregulated already at the 12h time point while other systems were expressed later only at the 24h time point, suggesting that Zorya responded the phage infection first. Zorya operates as a proton-driven motor that becomes activated after sensing of phage invasion, transferring the phage invasion signal through a cytoplasmic tail to recruit and activate the soluble effectors, which degrades the phage DNA^41^. Interestingly, Zorya has been found to interact with other phage defense systems such as Druantia and ietAS^42^. Our results thus suggest that co-expression of multiple defense system might have provided synergistic phage defense in *S. maltophilia*, or alternatively, expression of multiple defense systems could be result of population of cells activating different defense systems at different phases of phage replication cycle. In the future, it will be important to study if *sbb* mutations can potentiate higher levels of phage resistance by interacting with these phage defense systems proteins. Interestingly, while phage resistance trade-offs are often observed with pathogenic species^43^, including *S. maltophilia*^44^, we observed only a small growth cost of phage resistance in the lab conditions, even though the resistance was coupled with significant changes in the *S. maltophilia* secondary metabolism. Low pleiotropic cost could also explain why *S. maltophilia* was able to evolve phage resistance in the tomato rhizosphere and why phage resistant *S. maltophilia* mutants could suppress the pathogen *in planta*. Together, these results suggest that coevolutionary phage training could be used to directionally select for phage-resistant and more antagonistic *S. maltophilia* mutants through synergistic pleiotropy.

While we cannot fully explain how phage resistance mutations caused increase in *S. maltophilia* disease suppression, we observed comprehensive rewiring of *S. maltophilia* gene expression and metabolite production. In general, pathways associated with type IV secretion system were up-regulated in the presence of phage, which are vital for extracellular protein secretion and intraspecies antagonism^45^. Specifically, alkaline serine protease has been confirmed as one important mechanism for *S. maltophilia* to inhibit *R. solanacearum*^25^, whose biosynthesis process was rapidly activated by phage within 12 hours. We also found that phage infection rewired *S. maltophilia* secondary metabolite production, leading to secretion of several antimicrobial compounds including fumagillin, theophylline and α-ketoglutaric acid which have been reported to have health benefits in clinical context^46–48^. While we could establish link between metabolite production and gene expression for some of these compounds (theophylline, α-ketoglutaric acid, fumagillin), more future work is required to link these changes with observed mutations and activation of phage defense systems.

While phages are traditionally thought be specific to their host bacteria, this view has recently been challenged by studies showing that phages with broad host range are common in nature^49,50^, which could have community-wide effects in microbiomes. While the phage we used was highly specific to *S. maltophilia*, it could still indirectly modify the composition and diversity of rhizosphere bacterial community through interaction with its host bacterium. This finding is in line with previous studies where pathogen-specific phages were found to indirectly change the rhizosphere microbiome compostition^4^. In this study, we observed that the presence of *S. maltophilia* and its phage could prophylactically stabilize bacterial community composition and even increase bacterial community diversity when exposed to invasion by *R. solanacearum*, which alone clearly reduced the diversity and altered the composition of bacterial communities in line with a previous study^35^. These changes were likely due pathogen suppression and invasion limitation, as *R. solanacearum* densities were much lower in the presence of both *S. maltophilia* and its phage, resulting much lower disease progression. Interestingly, presence of phage did not have negative effect on *S. maltophilia* densities when measured at the end of the experiment (based on ASVs). Likely explanation for this is the evolution of *S. maltophilia* phage resistance, which was shown to emerge rapidly not only in the lab but also in the rhizosphere within 10 days.

At the level of co-occurrence networks, the presence of phage increased the number of species that were associated with *S. maltophilia* and led to exclusion of *R. solanacearum* from the major network module. As bacterial diversity has been considered vital for blocking pathogen invasion^51^, the pathogen suppression was also likely affected by the other bacterial species associated with *S. maltophilia* that also constrained the invasion of *R. solanacearum* in rhizosphere^52^. Interestingly, several of these bacterial species associated with the presence of *S. maltophilia* and its phage have previously been reported to be potential disease suppressive taxa^5,32–34^. Among these, we identified two key bacterial species, *E. ludwigii* YL-ENT31 and *P. baetica* PSE01, which exhibited enhanced antimicrobial activity against *R. solanacearum* in lab conditions. By conducting metabolomic analyses, we found that supernatant from *S. maltophilia*-phage coculture triggered increased production of several antimicrobial compounds by *E. ludwigii*, including ketoprofen acid, lactoferrin, dethiobiotin, and traumatic acid. Of these, ketoprofen has been identified as a quorum sensing inhibitor in *Pseudomonas aeruginosa*^53^, while lactoferrin has been reported to induce plant immune responses and to inhibit invasion of *R. solanacearum*^54^. Importantly, activation of bacterial pathogen suppressiveness by *S. maltophilia*-phage supernatant was not limited to these two bacterial strains but could be observed across several tomato rhizosphere bacterial taxa, indicating that increased disease suppressiveness was likely microbiome-wide phenomenon. Together these findings suggest that single *S. maltophilia*-specific phage can indirectly trigger microbiome-wide disease suppressiveness by activating bacterial antimicrobial activity. As *S. maltophilia*-specific phage had mainly positive indirect effects on bacterial diversity, the triggered pathogen suppression did not cause significant or long-lasting collateral damage to surrounding microbiota.

Manipulating microbial interactions for improved community function has been suggested a promising strategy to improve agricultural yields and control pathogens. Here we show that this could be achieved by using only two functionally different microbial species: plant growth-promoting bacterium and its phage. These findings are in contrast with often observed phage resistance trade-offs, highlighting the need to understand how non-pathogen species interact with their phages. In the future, it will be interesting to study interactions between other important plant growth-promoting bacteria and their phages to understand if our findings can be generalized across other bacterial and phage taxa. More work is also required to better understand the molecular basis of observed pleiotropic synergy between the phage resistance and antibiosis, and how plants perceive and adjust their gene expression in response to changes in microbial secondary metabolism. While the key focus of this study was on bacterial communities, metaviromics could help unravelling the cascading effects of *S. maltophilia* and its phage on rhizosphere phage communities and their host bacteria in the future. In conclusion, our findings provide a novel framework to improve pathogen biocontrol through phage-steering of bacterial community disease suppressiveness via activation of antimicrobial activity.

## Materials and methods

### Microbial strains and culture conditions

In this study, we used phage YL-STE-P and its *S. maltophilia* host bacterial strain YL-STE-01 as a model probiotic phage and bacterium pair. As a model pathogen, we used *R. pseudosolanacearum* QL-Rs1115 type strain, which has been sequenced and has been studied extensively (GenBank: GU390462)^55^. As the Ralstonia species classification based on genome information is relatively new, this strain is still referred as *R. solanacearum* throughout the text. As for model resident bacteria, we used *E. ludwigii* YL-ENT-31, which has previously been identified as a plant growth-promoting strain^3^ and *Pseudomonas baetica* PSE01 along with another 86 strains that were isolated from soil samples collected in a tomato field^3^ (Supplementary data S1, 118°57 E, 32°03’ N, Qilin, Nanjing, China). During the laboratory experiments, all bacteria were routinely cultured in a liquid nutrient medium (NB; 10.0 g of glucose, 5.0 g of peptone, 0.5 g of yeast extract, 3.0 g of beef extract, 1 L of H_2_O, pH 7.0) at 30 °C with shaking (170 rpm). The bacteria cells were harvested by centrifugation (6,000 rpm, 5 min), washed twice with sterile saline solution (0.9% NaCl), and diluted to a density of 10^6^-10^7^ colony-forming units (CFU) ml^−1^ based on the optical density 600 (OD_600_ = 0.5) before all follow-up experiments. To propagate the *S. maltophilia* phage YL-STE-P for the experiments, we co-cultured the host *S. maltophilia* bacterium and its phage in 3 ml of NB in six-well plates at 30 °C with shaking (170 rpm) with multiplicity of infection (MOI) of 10, where phages showed very high bacterial biomass reduction^3^. After 12h of co-culturing, bacterial cells were removed by centrifugation (12,000 rpm, 5 min), and supernatant was filtered through a 0.22-μm membrane filter to collect phage particles, which were stored at 4 °C until later use. Before all the experiments, phage culture was adjusted to 10^6^-10^7^ plaque-forming units (PFU) ml^−1^ by serial dilution in sterilized water.

### Quantifying the infectivity of YL-STE-P phage against other rhizosphere bacteria

We determined the infectivity range of *S. maltophilia* phage YL-STE-P against 88 phylogenetically distinct strains along with its natural host *S. maltophilia* YL-STE-01 and *R. pseudosolanacearum* QL-Rs1115 type strain. Phage resistance against YL-STE-P was quantified using 96-well microplates with 178 μL of liquid NB media at 30 °C with shaking (170 rpm). All wells were inoculated with 2 μL of bacteria and grown in the absence (20 μL of sterilized water, control) and the presence of YL-STE-P phage (20 μL of phage, MOI = 10) with eight biological replicates for each tested isolate. The biomass was examined after 24 hours of co-culturing as the OD_600_ using SpectraMax M5 Plate reader (Molecular Devices, Sunnyvale, CA). The phage resistance was calculated as the difference in bacterial growth in the absence (OD_600c_) and in the presence of phage (OD_600p_) treatments using the following formula: phage resistance = (OD_600p_ – OD_600c_)/OD_600c_. We also tested the growth curves of host *S. maltophilia* bacterium in the presence and absence of YL-STE-P phage with the same system by reading the OD_600_ every 4 hours from beginning to 32 hours and took final point read at 48 hours (Fig. 1a).

### Whole genome sequencing of bacteria and phage

To sequence the full genome of *S. maltophilia* YL-STE-01 and *E. ludwigii* YL-ENT-31bacteria and *S. maltophilia*-specific phage YL-STE-P, we first extracted bacterial DNA by using Bacteria DNA extraction Kit (OMEGA) and phage DNA with λ phage genomic DNA purification kit (ABigen) following manufacturer’s instructions. The DNA concentrations and quality were determined with NanoDrop 1000 spectrophotometer (Thermo Scientific) and electrophoresis for DNA shearing. High qualified DNA samples (OD_260_/OD_280_ = 1.8-2.0, > 6ug) were utilized to construct fragment library. All genomes were sequenced using a combination of PacBio and Illumina sequencing platforms (Shanghai BIOZERON Co., Ltd). The Illumina data was used to evaluate the complexity of the genome and correct the PacBio long reads. First, we used ABySS^56^ (v2.3.4) with multiple-Kmer parameters to achieve optimal results for genome assemblies. Secondly, canu^57^ (v2.2) was used to assemble the PacBio corrected long reads. Finally, GapCloser^58^ (v1.1.2) was applied to fill up the remaining local inner gaps and correct the single base polymorphism for the final assembly results. GeneMarkS^59^, tRNAscan-SE^60^ (v2.0.4) and RNAmmer^61^ (v1.2) were subsequently used to identify coding sequences, tRNA and rRNA regions, respectively. Moreover, we used eggNOG-mapper^62^ (v2.1.9) to annotate general gene functions and genomes were visualized using Circos^63^ (v0.64).

### Assessing the phage resistance and pathogen inhibition by evolved *S. maltophilia* colonies *in vitro*

To test how quickly *S. maltophilia* evolves resistance to its phage, the ancestral *S. maltophilia* bacterial cells were cultured in the absence (SM-Alone) and in the presence (SM+Phage) of its phage in 5 ml of NB medium (15-ml falcon tubes with loose lids to ensure aerobic growth conditions) at 30 °C with shaking (170 rpm). Both treatments were replicated for 10 times and cultures were established by first inoculating the replicate tubes with a final density of 10^6^ CFU/ml cells of isogenic *S. maltophilia* ancestral bacterium. Phage treatments were further inoculated with a final density of 10^7^ PFU/ml phage particles, resulting in MOI of 10. After co-culturing for 12 hours, we randomly selected 5 replicates from both treatments for destructive sampling and centrifuged (6,000 rpm, for 5 min) them to remove supernatant and frozen immediately in liquid-nitrogen to extract both DNA and RNA for re-sequencing and RNA-sequencing assay, respectively. After culturing for additional 24 hours, the remaining treatment replicates were homogenized and mixed together (50 μl, around 1% of total volume) to isolate evolved *S. maltophilia* strains and to collect DNA and RNA for sequencing. To isolate *S. maltophilia* colonies, culture suspensions were serially diluted and spread on nutrient agar plates (NA, 25 g l^−1^ agar with same nutrition content as NB media) and incubated at 30 °C for 24 hours. A total of 188 bacterial colonies (evolved mutants) were randomly selected and purified from phages by re-streaking on new agar plates. All evolved colonies from both treatments (evolved alone or in the presence of phages) were further examined for their phage resistance and inhibitory activity against *R. solanacearum* (as described earlier) and were classified as phage ‘resistant’ or ‘sensitive’ mutants’ based on phage resistance (> 5% and < -5%, respectively) as described earlier. Six representative colonies for both treatment groups were selected for genome re-sequencing.

### Genomic variation caused by *S. maltophilia* phage infection

We applied population re-sequencing method to identify mutations in evolved *S. maltophilia* host populations at 12h and 24h co-culture time points. In addition, eight *S. maltophilia* colonies evolved in the presence or absence of phage (N=4) were isolated and re-sequenced to determine mutations relative to ancestor *S. maltophilia* strain. Genomic DNA of each individual treatment replicate was extracted using a Bacteria DNA extraction Kit (OMEGA) following manufacturer’s instructions. High-quality DNA (> 3 µg, concentration > 30 ng/µl, OD_260_/OD_280_ = 1.80-2.00) was used for library construction and Illumina sequencing. Paired-end libraries (PE150) were sequenced by Illumina HiSeq PE (2 × 150) platform (Shanghai BIOZERON Co., Ltd). The raw pair-end reads were trimmed and quality controlled by Trimmomatic^64^ (v0.39) with default parameters. High quality reads were aligned to the reference genome (genome of *S. maltophilia* YL-STE-01) using BWA^65^ (r1188) with “bwa mem” mode. Variant calling was generated after quality control with parameters ‘QD < 2.0 || FS > 60.0 || MQ <40.0 || SOR > 10.0’ by GATK^66^ (v4.2.5.0) and filtered with VCFtools^67^ (v0.1.16) by parameters ‘--minQ 20 --minDP 4’. We used ANNOVAR^68^ (hg18) to detect genomic variants including SNPs and InDels. To measure the population differentiation, fixation index (Fst) was calculated also by VCFtools^67^ with default parameters.

### Transcriptional responses of *S. maltophilia* to phage infection

We applied transcriptomics to determine changes in *S. maltophilia* gene expression in response to phage infection at 12h and 24h co-culture time points. Total RNA was extracted from the cells using TRIzol® Reagent according the manufacturer’s instructions (Invitrogen) and genomic DNA was removed using DNase I (TaKara). High-quality RNA samples (> 10μg, OD_260_/OD_280_ = 1.8-2.2, OD_260_/OD_230_ ≥ 2.0, RIN ≥ 7, 28S:18S ≥ 1.0) were used to construct strand-specific libraries with TruSeq RNA sample preparation Kit from Illumina (San Diego, CA). Paired-end libraries were sequenced by Illumina NovaSeq 6000 sequencing platform (2 × 150, Shanghai BIOZERON Co., Ltd). The raw paired end reads were trimmed and quality controlled by Trimmomatic^64^ (v0.39) with parameters ‘SLIDINGWINDOW:4:15 MINLEN:75’ and then separately aligned to reference genome (genome of *S. maltophilia* YL-STE-01) with orientation mode using Rockhopper^69^. This software was also used to calculate gene counts with default parameters. We next used edgeR^70^ for differential expression analysis and clusterProfiler^71^ for gene set enrichment analysis (GSEA) based on KEGG pathways.

### Identification of key antimicrobial extracellular metabolites active against *R. solanacearum* using non-target metabolomics and pure matter validation for both *S. maltophilia* and *E. ludwigii*

To identify key antimicrobial extracellular metabolites inhibitory to *R. solanacearum*, we mixed 50 μL of *S. maltophilia* and *E. ludwigii* supernatant samples and 150 μL of 20 % acetonitrile methanol internal standard extractant by vortexing for 3 min followed by centrifugation (12,000 rpm, 4 °C, 10 min). Next, we transferred 150 μL of the supernatant and let it stand still at -20°C for 30 min before centrifugation (12,000 rpm, 4 °C, 3 min). The supernatants were collected for ultraperformance liquid chromatography/quadrupole time-of-flight mass spectrometry (UHPLC–QTOF–MS) at Shanghai Biotree Biotechnology, China. Mass spectrometry raw data were converted to the mzXML format using ProteoWizard^72^ (v3) and processed by R package XCMS^73^ (v3.2). We used CAMERA^74^ for peak annotation after XCMS data processing. *S. maltophilia* and *E. ludwigii* metabolite identification was performed using a combination of Human Metabolome (HMDB v5.0), MassBank of North America (MONA, https://massbank.us/), and Metlin (https://metlin.scripps.edu/) databases with R package MetaboAnalystR^75^. To examine the direct effect of fumagillin, theophylline, α-ketoglutaric acid, 1,2-Dioctanoyl PC (produced by *S. maltophilia*), ketoprofen acid, lactoferrin, dethiobiotin and traumatic acid (produced by *E. ludwigii*) on *R. solanacearum*, we cultured 2 μL (∼10^5^ CFU) of *R. solanacearum* with 178 μL of liquid NB media and 20 μL of sterilized water (control) or each above compounds (dissolved in sterilized water at a final concentration of 10 mM) with six biological replicates. The biomass was examined after 48 hours of culturing at 30 °C with shaking (170 rpm) as the optical density (OD_600_).

### Measuring the increase in *S. maltophilia* toxicity and alkali proteinase production by *S. maltophilia*-specific phage

The *S. maltophilia* was cultured in the presence or absence of its phage in 3 ml of NB medium and incubated at 30 °C temperatures with shaking at 170 rpm for 48 hours with 10 biological replicates and changes in bacterial biomass were measured as optical density (OD_600_). All cultures were centrifuged (12,000 rpm, 5 min) and filtered through 0.22-μm membrane filters to obtain supernatant without bacterial cells. Six out of ten replicates were used to measure the toxicity and alkali proteinase production as follows. We first collected 2 ml of cultures from treatments with phage and ultra-filtered them with 3 kDa membrane filters (4,000 rpm, 30 min). As phage particles were relatively larger, we collected the retentate as phage enrichment (Phage) and permeate as secondary metabolites (Metabolite, Supplementary Fig. S6a). To validate the absence of phage in the metabolite, we further used typical soft agar double-layer assay and found that no phage plaque formation was observed in the metabolite fraction. To measure the fraction toxicity to *S. maltophilia*, 2 μL of *S. maltophilia* were inoculated into 96-well microplates with 158 μL of liquid NB media and 20 μL of enriched phage (Phage), secondary metabolites (Metabolite) with 20 μL of sterilized water, a mix of 20 μL of enriched phage (Phage) and secondary metabolites (Metabolite), or 40 μL of sterilized water (Water). All treatments were grown for 24 hours at 30°C with shaking (170 rpm) before measuring bacterial densities as optical density (OD_600_). The remaining supernatants cultured in the absence (SM-Alone) or presence of phage (SM+Phage) were further used to measure the alkali proteinase productivity with alkaline proteinase (AKP) activity assay kit (Sangon Biotech, Shanghai, China) following the manufacturer’s instructions. Another four supernatant replicates cultured in the in the absence (SM-Alone) or presence (SM+Phage) of phage were used for the non-targeted metabolomics assay.

### The experimental setup of greenhouse experiments and collection of soil samples

We used a commercially available ‘Red Dwarf’ tomato cultivar (*Lycopersicon esculentum*, Shouguangxuran Agricultural Technology Co., Ltd.) as the model plant^3^, which is highly susceptible to *R. solanacearum*. At the three-leaf stage, tomato plants cultured on aseptic commercial substrate were transplanted into 6-plant cell trays before starting the experiments. Each cell contained 200 g of thoroughly mixed topsoil, which was collected from the tomato field in Qilin town, Nanjing, China^3^. The soil is typical yellow-brown soil, containing 24.0 gkg^−1^ of organic matter, 1.7 g kg^−1^ of total nitrogen, 173.1 mg kg^−1^ of available phosphorus, and 178 mg kg^−1^ of available potassium with pH of 5.8^5^. After removing surface debris on the ground, soil was collected from a depth of 5-20 cm. All experimental soils were sieved (<4 mm) and homogenized thoroughly before use. To prepare sterilized soils for some of the experiment, a subset of homogenized soils was sterilized with gamma irradiation at 75kGy. All the plants were grown in same conditions in a glasshouse facility with a natural temperature variation ranging between 28 and 35 °C. The *S. maltophilia* was inoculated to plant roots at a final concentration of 10^5^-10^6^ CFU g^−1^ soil and its phage at the same time at a final concentration of 10^6^-10^7^ PFU g^−1^ soil (M.O.I = 10). Plants in treatments without inoculated strains were inoculated with the same amount of sterile water. Three separate greenhouse experiments were conducted using this setup, which are described in more detail below.

#### Experiment 1: Comparing the disease suppressiveness of phage resistant S. maltophilia mutants

A total of 240 plants were used to quantify the disease suppressiveness of lab-evolved colonies where each 6-well tray was seen as an individual biological replicate (four replicates per treatment in total). Plants were separated into 10 treatments with different inoculations including one ancestor *S. maltophilia*, four *S. maltophilia* colonies isolated from ‘Evolved with phage’ treatment, four *S. maltophilia* colonies from ‘Evolved alone’ treatments in previous lab-evolutionary assay, and same amount of water as control. All plants were also inoculated with *R. solanacearum* QL-Rs1115 type strain. All bacterial colonies were inoculated at the same time at a final concentration of 10^5^-10^6^ CFU g^−1^ soil. The experiment was finished 30 d after the bacterial inoculations when the disease progression had reached a stable level. The plant disease severity was quantified during the experiment based on disease index^76^, which ranged from 0 to 4 (0, no signs of wilting; 1, 1%-25% leaf area wilted; 2, 26%-50% leaf area wilted; 3, 51%-75% leaf area wilted; and 4, 76%-100% leaf area wilted)^77^ and finally normalized as percentage of diseased plants per replicate and treatment.

#### Experiment 2: determining S. maltophilia phage resistance evolution in the rhizosphere

Another 8 plants growing in sterilized soils were separated into two treatments (four each) where ancestral *S. maltophilia* colonies were grown in the presence and absence of phage (N=4). The ancestral *S. maltophilia* colony was inoculated to each plant rhizosphere at a final concentration of 10^5^-10^6^ CFU g^−1^ soil and four plants in phage treatment were also inoculated with *S. maltophilia* phage at a final concentration of 10^6^-10^7^ PFU g^−1^ soil (phage treatment). We considered each plant as an individual selection line and collected rhizosphere soil samples from each plant after 10 days of inoculation. One evolved *S. maltophilia* colony per treatment replicate was isolated and purified from phages with culturing on with nutrient medium plates as described previousluy^3^ and the taxonomic identity of colonies were verified using 16S rRNA sequencing. All selected colonies were full genome re-sequenced and quantified regarding the antibiotic activity against *R. solanacearum* and phage resistance against ancestral phage as described previously.

#### Experiment 3: Assessing the effect of S. maltophilia-phage evolution on rhizosphere microbiome

We used the scaled-down “rhizobox” system which consists of central growth section lined up with soil-containing individual nylon bags that can be removed during experiments, allowing repeated, non-destructive sampling of a subset of rhizosphere without damaging plant roots in this experiment^23^. A total of 72 plants evenly divided into three treatments with four replicates (‘Rs-Control’, ‘SM-Alone’ and ‘SM+Phage’, each included six plants) were used to determine the role of *S. maltophilia* and its phage on microbiota diversity, community composition and co-occurrence networks. After 10 d from inoculation of *S. maltophilia* bacterium and phage, we removed one rhizo-bag from each plant to collect rhizosphere samples which were classified as the ‘pre-infection stage’ samples. Soils collected from each 6-cell tray were mixed thoroughly within treatments and stored at -80°C immediately. Next, we inoculated *R. solanacearum* QL-Rs1115 type strain at a final concentration of 10^5^-10^6^ CFU g^−1^ soil. After 20 days, another rhizo-bag from each plant was also removed (‘post-infection stage’ sample) and processed similar to ‘initial stage’ samples to collect information on pathogen density (qPCR; methodsdescribed in the next section) and rhizosphere microbiome (16S rRNA sequencing; method described in the next section). Finally, we quantified the plant disease severity with same criteria as described previously.

### Soil sample preparation, qPCR assay and 16S rRNA amplicon sequencing during the experiment 3

The total microbial DNA from all collected soil samples from both sampling times was extracted using the Power Soil DNA Isolation kit (Mo Bio Laboratories) according to the manufacturer’s protocols. The extracted DNA was first used to determine the pathogen density by using qPCR targeting the *R. solanacearum*-specific *fliC* gene as copy numbers per gram of soil^4^. Second, we used 16S rRNA amplicon sequencing to determine bacterial community composition and diversity in the same samples. The V4-V5 region of the bacterial 16S rRNA genes was amplified using primers 515F 5’-barcode-GTGCCAGCMGCCGCGG)-3’ and 907R 5’-CCGTCAATTCMTTTRAGTTT-3’ where barcodes included an eight-base sequence unique to each sample. The amplicon library was paired-end sequenced (2 × 250) on an Illumina MiSeq platform (Shanghai BIOZERON Co., Ltd) by following standard protocols. Sequence reads that passed standard quality-control^76^ were dereplicated and subjected to the DADA2^78^ algorithm by QIIME 2^79^ (v2022.8). The trimming and filtering were performed on paired reads with a maximum of two expected errors per read (maxEE = 2). After merging the paired reads and chimera filtering, the phylogenetic affiliation of each 16S rRNA gene sequence (herein called ASVs) was analyzed using RDP Classifier^80^ (2.10.2) against the silva (SSU132) 16S rRNA database^81^ with confidence threshold levels of 80%. This resulted in a dataset comprising 912,825 reads and 8,126 ASVs. To match the ASVs with our focal bacterial strains, the 16S sequences of *S. maltophilia* YL-STE-01, *E. ludwigii* YL-ENT-31, *R. solanacearum* QL-Rs1115 and type strains downloaded from NCBI database were collected and aligned with representative ASV sequences using MUSCLE^82^ (UPGMB method) algorithm. The alignment was further visualized as phylogenetic trees with MEGA-X^83^ (v10.2.2) using the neighbour-joining method with bootstrapping (999 replicates).

### Assessing the impact of *S. maltophilia* and its phage on the tomato rhizosphere microbiota during the experiment 3

To determine the influence of *S. maltophilia*–phage co-evolution during pathogen invasion at community level, we calculated the Shannon diversity for each sample and estimated the fold change between ‘final stage’ and ‘initial’ stage. For the effect on individual resident bacteria, we quantified the effect of *R. solanacearum* invasion by calculating the changes of ASVs relative abundances between final stage and initial stage. Next, we measured the effect of *S. maltophilia*–phage on these taxa abundance changes using the following formula: Effect = (Change_SM+Phage_ – Change_SM-Phage_)/Average abundance. In addition, to reveal the impact on potential microbial interactions, we selected a subset of major ASVs occurring in at least half of samples with average proportion > 0.02%, resulting in a total of 470 ASVs. Subsequently, we quantified their pairwise interaction potential with MIC algorithm (R package minerva^84^ v1.4.3) with a combination of pre-infection and post-infection stage data. Those connections with MIC value > 0.8 were kept and constructed for networks. We used ggClusterNet^85^ (v0.1.0) to calculate the network properties and visualized them by Gephi^86^ (0.10.1).

### Determining the cascading effect of *S. maltophilia*–specific phage on the suppressiveness of resident bacteria against *R. solanacearum*

To assess how *S. maltophilia*–specific phage interaction affects resident bacteria, we used the 88 bacterial tomato rhizosphere isolates which were resistant to YL-STE-01P phage (which included two candidate species, *E. ludwigii* YL-ENT-31 and *P. baetica* PSE1) to determine the effect of *S. maltophilia* phage on pathogen-resident bacteria interactions. As the *S. maltophilia* phage cannot infect either *E. ludwigii* or *P. baetica* (or other 86 bacterial species), we first explored the pairwise microbial interactions between *S. maltophilia* and *E. ludwigii* or *P. baetica* mediated by secondary metabolites. To obtain their extracellular metabolites, 2 μL of *E. ludwigii* or *P. baetica* (∼10^5^ CFU) was cultured with 178 μL of liquid NB media and 20 μL of sterilized water (*E. ludwigii*^S^ or *P. baetica*^S^), while 2 μL of *S. maltophilia* (∼10^5^ CFU) was cultured with 178 μL of liquid NB media in the absence (20 μL of sterilized water, SM-Alone^S^) and presence of *S. maltophilia* phage (20 μL of phage, MOI = 10, SM+Phage^S^) with eight biological replicates each. After 48 hours of culturing at 30 °C with shaking (170 rpm), we centrifuged all treatments (12,000 rpm, 5 min) and filtered them through a 0.22-μm membrane filter to harvest supernatants. To determine the metabolite-mediated pairwise interactions, 2 μL of target bacterium (*S. maltophilia*, *E. ludwigii* or *P. baetica*, ∼10^5^ CFU) was treated with 20 μL of supernatant (*E. ludwigii*^S^, *P. baetica*^S^, SM-Alone^S^ or SM+Phage^S^, sterilized water as control) and 178 μL of liquid NB media with eight replicates. The biomass was examined after 24 hours of culturing at 30 °C with shaking (170 rpm) as the OD_600_.

To measure the indirect effect of *S. maltophilia*–specific phage on the suppressiveness of remaining 86 resident bacteria (including both *E. ludwigii* and *P. baetica*) against *R. solanacearum*, we cultured 2 μL (∼10^5^ CFU) of bacterium (bacterium^S^) and sterilized water (Water) with 178 μL of liquid NB media and 20 μL of previously collected SM-Alone^S^, SM+Phage^S^ supernatants or sterilized water with six replicates at 30 °C with shaking (170 rpm). After 48 hours of culturing, we centrifuged all treatments (12,000 rpm, 5 min) and filtered them through 0.22-μm membrane filters to obtain supernatants. The inhibition activity of each supernatant was then determined by co-culturing with *R. solanacearum* following the same system we used for *S. maltophilia*. In addition, four replications of *E. ludwigii* supernatants cultured with SM-Alone^S^ and SM+Phage^S^ were collected for non-target metabolomic assay to identify key compounds enriched by *S. maltophilia*–specific phage. We used iTols^87^ (v6) to visualize the phylogenetic relationships of resident bacteria along with the effect of *S. maltophilia*-specific phage on their inhibition activity to *R. solanacearum*.

### Statistical analysis

All downstream statistical analyses were conducted using R platform^88^ (v 4.3.1). We used the unpaired t-test to compare the differences between two treatments in both microbiome analysis and culture-dependent experiments. In case of unbalanced data, the unpaired Wilcoxon test was used. For comparison among multiple treatments, the Tukey’s multiple comparisons were used after one-way ANOVA. Differences in community and metabolome composition were compared using the PERMANOVA test. The Shannon index was calculated using the ‘diversity’ function in the vegan package based on the taxa abundance matrix. Principal components analysis (PCA) and principal-coordinates analysis (PCoA) based on Bray-Curtis distances were performed by stats and ape packages, respectively. For transcriptomic data, the edgeR^70^ package was used for differential expression analysis. For metabolomic data, the ropls package was used to calculate VIP value for differential analysis after orthogonal partial least squares discriminant analysis (OPLS-DA). *P*-values <0.05 were considered statistically significant.

## Data availability

All sequencing data generated for this manuscript is available in National Genomics Data Center and can be accessed using BioProject: PRJCA052801.

## Acknowledgements

This study was financially supported by the National Natural Science Foundation of China (42325704, 42407171, 42090064 and 42377118), the Natural Science Foundation of Jiangsu Province (BK20240194), China Postdoctoral Science Foundation (GZB20240311, 2025T180072, 2024M760612) and Research Council of Finland (project 355505). This study was also supported by the high-performance computing public platform of Nanjing Agricultural University.

## References

1 Wheatley, R. M., Holtappels, D. & Koskella, B. Evaluation of bacteriophages as a signature of microbiome health: a systematic review and meta-analysis. The Lancet Microbe (2025). 10.1016/j.lanmic.2025.101196

2 Strathdee, S. A., Hatfull, G. F., Mutalik, V. K. & Schooley, R. T. Phage therapy: From biological mechanisms to future directions. Cell 186, 17–31 (2023). 10.1016/j.cell.2022.11.017

3 Yang, K. et al. Rhizosphere phage communities drive soil suppressiveness to bacterial wilt disease. Microbiome 11, 16 (2023). 10.1186/s40168-023-01463-8

4 Wang, X. et al. Phage combination therapies for bacterial wilt disease in tomato. Nature Biotechnology 37, 1513–1520 (2019). 10.1038/s41587-019-0328-3

5 Wang, X. et al. Phages enhance both phytopathogen density control and rhizosphere microbiome suppressiveness. mBio 15, e03016–03023 (2024). 10.1128/mbio.03016-23

6 Hsu, B. B. et al. Dynamic Modulation of the Gut Microbiota and Metabolome by Bacteriophages in a Mouse Model. Cell Host Microbe 25, 803–814 e805 (2019). 10.1016/j.chom.2019.05.001

7 Liang, X. et al. Bacteriophage-driven microbial phenotypic heterogeneity: ecological and biogeochemical importance. NPJ Biofilms Microbiomes 11, 82 (2025). 10.1038/s41522-025-00727-5

8 Wang, J. et al. Phage selection drives resistance-virulence trade-offs in Ralstonia solanacearum plant-pathogenic bacterium irrespective of the growth temperature. Evolution Letters 8, 253–266 (2024). 10.1093/evlett/qrad056

9 Rabsch, W. et al. FepA- and TonB-Dependent Bacteriophage H8: Receptor Binding and Genomic Sequence. Journal of Bacteriology 189, 5658–5674 (2007). 10.1128/jb.00437-07

10 Burmeister, A. R. & Turner, P. E. Trading-off and trading-up in the world of bacteria-phage evolution. Curr Biol 30, R1120–R1124 (2020). 10.1016/j.cub.2020.07.036

11 Burmeister, A. R. et al. Pleiotropy complicates a trade-off between phage resistance and antibiotic resistance. Proceedings of the National Academy of Sciences 117, 11207–11216 (2020). 10.1073/pnas.1919888117

12 Kronheim, S. et al. A chemical defence against phage infection. Nature 564, 283–286 (2018). 10.1038/s41586-018-0767-x

13 Bernheim, A. et al. Prokaryotic viperins produce diverse antiviral molecules. Nature 589, 120–124 (2021). 10.1038/s41586-020-2762-2

14 Zang, Z. et al. Streptomyces secretes a siderophore that sensitizes competitor bacteria to phage infection. Nature Microbiology 10, 362–373 (2025). 10.1038/s41564-024-01910-8

15 Brady, T. S. et al. Bystander Phage Therapy: Inducing Host-Associated Bacteria to Produce Antimicrobial Toxins against the Pathogen Using Phages. Antibiotics (Basel) 7 (2018). 10.3390/antibiotics7040105

16 An, S.-q. & Berg, G. Stenotrophomonas maltophilia. Trends in Microbiology 26, 637–638 (2018). 10.1016/j.tim.2018.04.006

17 Vailleau, F. & Genin, S. Ralstonia solanacearum: An Arsenal of Virulence Strategies and Prospects for Resistance. Annual Review of Phytopathology 61, 25–47 (2023). 10.1146/annurev-phyto-021622-104551

18 Wang, Q. et al. LotS/LotR/Clp, a novel signal pathway responding to temperature, modulating protease expression via c-di-GMP mediated manner in Stenotrophomonas maltophilia FF11. Microbiological Research 214, 60–73 (2018). 10.1016/j.micres.2018.05.014

19 Shereda, R. D. et al. Ssb as an Organizer/mobilizer of Genome Maintenance Complexes. Critical Reviews in Biochemistry and Molecular Biology (2008).

20 Han, P. et al. Characterization of the Bacteriophage BUCT603 and Therapeutic Potential Evaluation Against Drug-Resistant Stenotrophomonas maltophilia in a Mouse Model. Frontiers in Microbiology **Volume** 13 **- 2022** (2022).

21 Tesson, F. et al. Systematic and quantitative view of the antiviral arsenal of prokaryotes. Nature Communications 13, 2561 (2022). 10.1038/s41467-022-30269-9

22 Payne, L. J. et al. Identification and classification of antiviral defence systems in bacteria and archaea with PADLOC reveals new system types. Nucleic Acids Research 49, 10868–10878 (2021). 10.1093/nar/gkab883

23 Wei, Z. et al. Initial soil microbiome composition and functioning predetermine future plant health. Science Advances 5, eaaw0759 (2019). 10.1126/sciadv.aaw0759

24 Windhorst, S. et al. The major extracellular protease of the nosocomial pathogen Stenotrophomonas maltophilia: characterization of the protein and molecular cloning of the gene. J Biol Chem 277, 11042–11049 (2002). 10.1074/jbc.M109525200

25 Elhalag, K. M., Messiha, N. A., Emara, H. M. & Abdallah, S. A. Evaluation of antibacterial activity of Stenotrophomonas maltophilia against Ralstonia solanacearum under different application conditions. J Appl Microbiol 120, 1629–1645 (2016). 10.1111/jam.13097

26 Krissinel, E. & Henrick, K. Inference of Macromolecular Assemblies from Crystalline State. Journal of Molecular Biology 372, 774–797 (2007). 10.1016/j.jmb.2007.05.022

27 Xue, L. C., Rodrigues, J. P., Kastritis, P. L., Bonvin, A. M. & Vangone, A. PRODIGY: a web server for predicting the binding affinity of protein–protein complexes. Bioinformatics 32, 3676–3678 (2016). 10.1093/bioinformatics/btw514

28 Passaro, S. et al. Boltz-2: Towards Accurate and Efficient Binding Affinity Prediction. bioRxiv, 2025.2006.2014.659707 (2025). 10.1101/2025.06.14.659707

29 Ahad, A. et al. A green marriage: the union of theophylline’s catalytic activity and healing potential. RSC Adv 15, 18338–18357 (2025). 10.1039/d4ra08479a

30 Chen, X., Dong, X., Liu, J., Luo, Q. & Liu, L. Pathway engineering of Escherichia coli for α-ketoglutaric acid production. Biotechnology and Bioengineering 117, 2791–2801 (2020). 10.1002/bit.27456

31 Griffith, E. C. et al. Molecular recognition of angiogenesis inhibitors fumagillin and ovalicin by methionine aminopeptidase 2. Proceedings of the National Academy of Sciences 95, 15183–15188 (1998). 10.1073/pnas.95.26.15183

32 Li, C. et al. Massilia rhizosphaerae sp. nov., a rice-associated rhizobacterium with antibacterial activity against Ralstonia solanacearum. Int J Syst Evol Microbiol 71 (2021). 10.1099/ijsem.0.005009

33 Karthikeyan, A., Kanchanadevi, K. & Nicodemus, A. Effect of Frankia and Micromonospora on growth and health improvement in Casuarina clones. Journal of Forest Research 27, 128–132 (2022).

34 Matsumoto, H. et al. Bacterial seed endophyte shapes disease resistance in rice. Nat Plants 7, 60–72 (2021). 10.1038/s41477-020-00826-5

35 Wei, Z. et al. Ralstonia solanacearum pathogen disrupts bacterial rhizosphere microbiome during an invasion. Soil Biology and Biochemistry 118, 8–17 (2018). 10.1016/j.soilbio.2017.11.012

36 Neshat, M. et al. Canola inoculation with Pseudomonas baetica R27N3 under salt stress condition improved antioxidant defense and increased expression of salt resistance elements. Industrial Crops and Products 206, 117648 (2023). 10.1016/j.indcrop.2023.117648

37 Gu, S. et al. Competition for iron drives phytopathogen control by natural rhizosphere microbiomes. Nature Microbiology 5, 1002–1010 (2020). 10.1038/s41564-020-0719-8

38 Kell, D. B., Heyden, E. L. & Pretorius, E. The Biology of Lactoferrin, an Iron-Binding Protein That Can Help Defend Against Viruses and Bacteria. Frontiers in Immunology **Volume** 11 - 2020 (2020).

39 Advocating for phage therapy. Nature Microbiology 9, 1397–1398 (2024). 10.1038/s41564-024-01733-7

40 Jdeed, G., Morozova, V. V. & Tikunova, N. V. Genome Analysis of Anti-Phage Defense Systems and Defense Islands in Stenotrophomonas maltophilia: Preservation and Variability. Viruses 16 (2024).

41 Hu, H. et al. Structure and mechanism of the Zorya anti-phage defence system. Nature 639, 1093–1101 (2025). 10.1038/s41586-024-08493-8

42 Wu, Y. et al. Bacterial defense systems exhibit synergistic anti-phage activity. Cell Host & Microbe 32, 557–572.e556 (2024). 10.1016/j.chom.2024.01.015

43 Mangalea, M. R. & Duerkop, B. A. Fitness Trade-Offs Resulting from Bacteriophage Resistance Potentiate Synergistic Antibacterial Strategies. Infect Immun 88 (2020). 10.1128/IAI.00926-19

44 Han, P., Lin, W., Fan, H. & Tong, Y. Characterization of phage evolution and phage resistance in drug-resistant Stenotrophomonas maltophilia. Journal of Virology 98, e01249–01223 (2024). 10.1128/jvi.01249-23

45 Bayer-Santos, E. et al. The opportunistic pathogen Stenotrophomonas maltophilia utilizes a type IV secretion system for interbacterial killing. PLOS Pathogens 15, e1007651 (2019). 10.1371/journal.ppat.1007651

46 Asadi Shahmirzadi, A., et al. Alpha-Ketoglutarate, an Endogenous Metabolite, Extends Lifespan and Compresses Morbidity in Aging Mice. Cell Metab 32, 447–456 e446 (2020). 10.1016/j.cmet.2020.08.004

47 Molina, J. M. et al. Fumagillin treatment of intestinal microsporidiosis. N Engl J Med 346, 1963–1969 (2002). 10.1056/NEJMoa012924

48 Larsen, S. D. et al. Synthesis and Biological Activity of Analogues of the Antidiabetic/Antiobesity Agent 3-Guanidinopropionic Acid:[ Discovery of a Novel Aminoguanidinoacetic Acid Antidiabetic Agent. Journal of Medicinal Chemistry 44, 1217–1230 (2001). 10.1021/jm000095f

49 Bignaud, A. et al. Phages with a broad host range are common across ecosystems. Nature Microbiology (2025). 10.1038/s41564-025-02108-2

50 Wirbel, J. et al. Long-read metagenomics reveals phage dynamics in the human gut microbiome. Nature (2025). 10.1038/s41586-025-09786-2

51 Spragge, F. et al. Microbiome diversity protects against pathogens by nutrient blocking. Science 382, eadj3502 10.1126/science.adj3502

52 Li, M. et al. Indirect reduction of Ralstonia solanacearum via pathogen helper inhibition. The ISME Journal 16, 868–875 (2022). 10.1038/s41396-021-01126-2

53 Mirpour, M. & Zahmatkesh, H. Ketoprofen attenuates Las/Rhl quorum-sensing (QS) systems of Pseudomonas aeruginosa: molecular and docking studies. Mol Biol Rep 51, 133 (2024). 10.1007/s11033-023-09071-3

54 Buziashvili, A. & Yemets, A. Lactoferrin and its role in biotechnological strategies for plant defense against pathogens. Transgenic Res 32, 1–16 (2023). 10.1007/s11248-022-00331-9

55 Tan, S. et al. Bacillus amyloliquefaciens T-5 may prevent Ralstonia solanacearum infection through competitive exclusion. Biology and Fertility of Soils 52, 341–351 (2016). 10.1007/s00374-015-1079-z

56 Jackman, S. D. et al. ABySS 2.0: resource-efficient assembly of large genomes using a Bloom filter. Genome Research 27, 768–777 (2017).

57 Koren, S. et al. Canu: scalable and accurate long-read assembly via adaptive k-mer weighting and repeat separation. Genome Research 27, 722–736 (2017).

58 Luo, R. et al. SOAPdenovo2: an empirically improved memory-efficient short-read de novo assembler. GigaScience 1, 2047–2217X-2041-2018 (2012). 10.1186/2047-217X-1-18

59 Besemer, J., Lomsadze, A. & Borodovsky, M. GeneMarkS: a self-training method for prediction of gene starts in microbial genomes. Implications for finding sequence motifs in regulatory regions. Nucleic Acids Research 29, 2607–2618 (2001). 10.1093/nar/29.12.2607

60 Chan, Patricia P., Lin, Brian Y., Mak, Allysia J. & Lowe, Todd M. tRNAscan-SE 2.0: improved detection and functional classification of transfer RNA genes. Nucleic Acids Research 49, 9077–9096 (2021). 10.1093/nar/gkab688

61 Lagesen, K. et al. RNAmmer: consistent and rapid annotation of ribosomal RNA genes. Nucleic Acids Research 35, 3100–3108 (2007). 10.1093/nar/gkm160

62 Cantalapiedra, C. P., Hernández-Plaza, A., Letunic, I., Bork, P. & Huerta-Cepas, J. eggNOG-mapper v2: Functional Annotation, Orthology Assignments, and Domain Prediction at the Metagenomic Scale. Molecular Biology and Evolution 38, 5825–5829 (2021). 10.1093/molbev/msab293

63 Krzywinski, M. I. et al. Circos: An information aesthetic for comparative genomics. Genome Research (2009).

64 Bolger, A. M., Lohse, M. & Usadel, B. Trimmomatic: a flexible trimmer for Illumina sequence data. Bioinformatics 30, 2114–2120 (2014). 10.1093/bioinformatics/btu170

65 Li, H. & Durbin, R. Fast and accurate long-read alignment with Burrows–Wheeler transform. Bioinformatics 26, 589–595 (2010). 10.1093/bioinformatics/btp698

66 Morris, J. A., Randall, J., Maller, J. B. & Barrett, J.

67 Danecek, P. et al. The variant call format and VCFtools. Bioinformatics 27, 2156–2158 (2011). 10.1093/bioinformatics/btr330

68 Wang, K., Li, M. & Hakonarson, H. ANNOVAR: functional annotation of genetic variants from high-throughput sequencing data. Nucleic Acids Research 38, e164–e164 (2010). 10.1093/nar/gkq603

69 Tjaden, B. De novo assembly of bacterial transcriptomes from RNA-seq data. Genome Biology 16, 1 (2015). 10.1186/s13059-014-0572-2

70 Robinson, M. D., McCarthy, D. J. & Smyth, G. K. edgeR: a Bioconductor package for differential expression analysis of digital gene expression data. Bioinformatics 26, 139–140 (2010). 10.1093/bioinformatics/btp616

71 Wu, T. et al. clusterProfiler 4.0: A universal enrichment tool for interpreting omics data. The Innovation 2, 100141 (2021). 10.1016/j.xinn.2021.100141

72 Chambers, M. C. et al. A cross-platform toolkit for mass spectrometry and proteomics. Nature Biotechnology 30, 918–920 (2012). 10.1038/nbt.2377

73 Benton, H. P., Wong, D. M., Trauger, S. A. & Siuzdak, G. XCMS2: Processing Tandem Mass Spectrometry Data for Metabolite Identification and Structural Characterization. Analytical Chemistry 80, 6382–6389 (2008). 10.1021/ac800795f

74 Kuhl, C., Tautenhahn, R., Böttcher, C., Larson, T. R. & Neumann, S. CAMERA: An Integrated Strategy for Compound Spectra Extraction and Annotation of Liquid Chromatography/Mass Spectrometry Data Sets. Analytical Chemistry 84, 283–289 (2012). 10.1021/ac202450g

75 Pang, Z., Chong, J., Li, S. & Xia, J. MetaboAnalystR 3.0: Toward an Optimized Workflow for Global Metabolomics. Metabolites 10 (2020).

76 Yang, K. et al. RIN enhances plant disease resistance via root exudate-mediated assembly of disease-suppressive rhizosphere microbiota. Molecular Plant 16, 1379–1395 (2023). 10.1016/j.molp.2023.08.004

77 Tans-Kersten, J., Brown, D. & Allen, C. Swimming Motility, a Virulence Trait of Ralstonia solanacearum, Is Regulated by FlhDC and the Plant Host Environment. Molecular Plant-Microbe Interactions® 17, 686–695 (2004). 10.1094/MPMI.2004.17.6.686

78 Callahan, B. J. et al. DADA2: High-resolution sample inference from Illumina amplicon data. Nature Methods 13, 581–583 (2016). 10.1038/nmeth.3869

79 Bolyen, E. et al. Reproducible, interactive, scalable and extensible microbiome data science using QIIME 2. Nature Biotechnology 37, 852–857 (2019). 10.1038/s41587-019-0209-9

80 Wang, Q., Garrity George, M., Tiedje James, M. & Cole James, R. Naïve Bayesian Classifier for Rapid Assignment of rRNA Sequences into the New Bacterial Taxonomy. Applied and Environmental Microbiology 73, 5261–5267 (2007). 10.1128/AEM.00062-07

81 Quast, C. et al. The SILVA ribosomal RNA gene database project: improved data processing and web-based tools. Nucleic Acids Research 41, D590–D596 (2013). 10.1093/nar/gks1219

82 Edgar, R. C. MUSCLE: multiple sequence alignment with high accuracy and high throughput. Nucleic acids research 32 **5**, 1792–1797 (2004).

83 Kumar, S., Stecher, G., Li, M., Knyaz, C. & Tamura, K. MEGA X: Molecular Evolutionary Genetics Analysis across Computing Platforms. Molecular Biology and Evolution 35, 1547–1549 (2018). 10.1093/molbev/msy096

84 Albanese, D. et al. minerva and minepy: a C engine for the MINE suite and its R, Python and MATLAB wrappers. Bioinformatics 29, 407–408 (2013). 10.1093/bioinformatics/bts707

85 Wen, T. et al. ggClusterNet: An R package for microbiome network analysis and modularity-based multiple network layouts. iMeta 1, e32 (2022). 10.1002/imt2.32

86 Bastian, M., Heymann, S. & Jacomy, M. Gephi: An Open Source Software for Exploring and Manipulating Networks. Proceedings of the International AAAI Conference on Web and Social Media (2009).

87 Letunic, I. & Bork, P. Interactive Tree of Life (iTOL) v6: recent updates to the phylogenetic tree display and annotation tool. Nucleic Acids Research 52, W78–W82 (2024). 10.1093/nar/gkae268

88 Team, R. C. R: A Language and Environment for Statistical Computing. (2023).

