## Supplemental Figure 1-8; Supplemental Table 1-3 for "Probiotic phage steering of disease suppressive rhizosphere microbiome"

**Supplementary Figure 1**

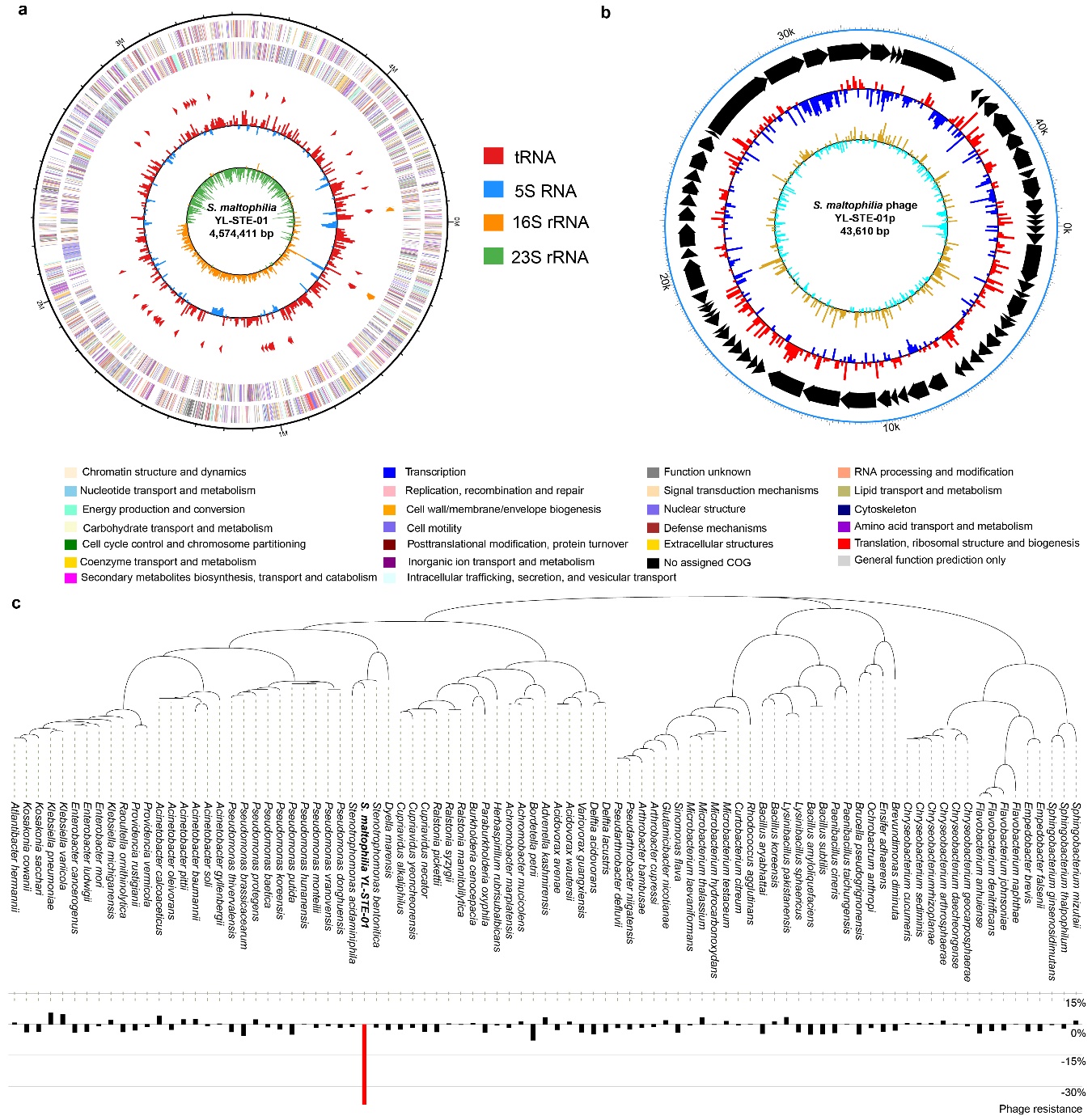

**Supplementary Figure 1. The full genomes of *S. maltophilia* YL-STE-01 and phage YL-STE-01p as well as the host range of *S. maltophilia* phage across 88 tomato rhizophere bacterial isolates.** **a-b**, The genome map of *S. maltophilia* YL-STE-01 and phage YL-STE-01p. **c**, The host specificity of phage YL-STE-01p based on the relative phage resistance on 88 different bacterial species across four phyla (black and red denotes for high and low phage resistance). The phylogenetic tree was constructed by 16s sequences using neighbor-joining method.

**Supplementary Figure 2**

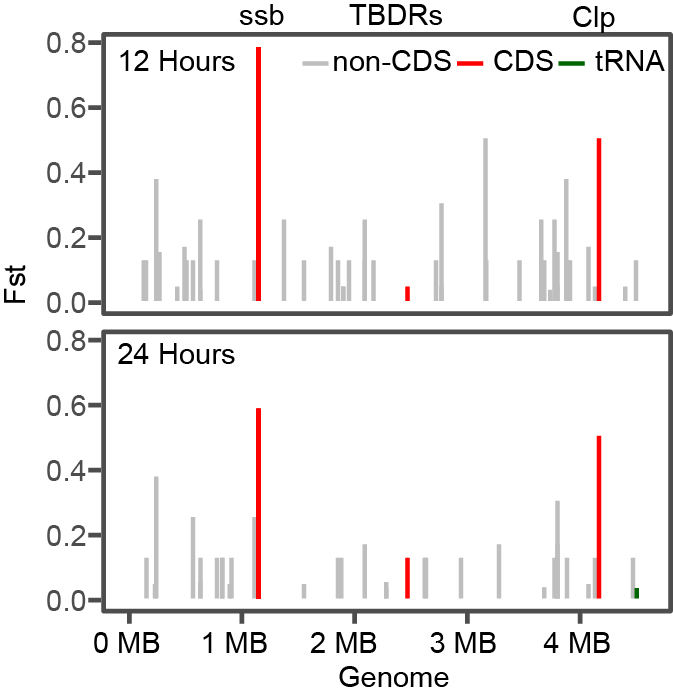

**Supplementary Figure 2. Fixation index (Fst) of observed genomic variants in *S. maltophilia* genome in the presence of phage at 12 (top panel) and 24 (bottom panel) hour sampling time points based on population re-sequencing (*n* = 5)**. Line colors denote for different genomic regions including (non-) coding sequences (CDS) and tRNA. The nonsynonymous mutations observed at both 12 and 24 hours are labeled with the annotated gene functions.

**Supplementary Figure 3**

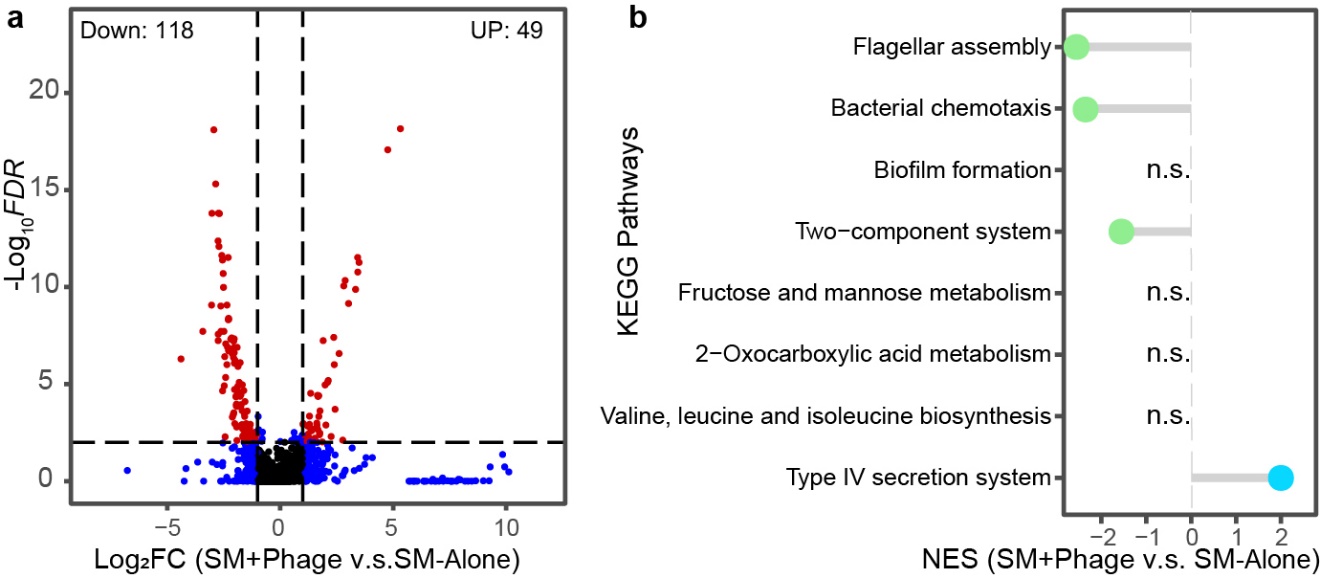

**Supplementary Figure 3.** **Changes in S*. maltophilia* gene expression at 12-hour sampling time point in the presence of phage. a,** Differentially expressed genes in the presence of phage. Red points denote significantly up or down regulated genes with *FDR* < 0.01 and |Log_2_ Fold Change| >1 (DEseq, *n* = 5); a total of 49 genes were up-regulated while 118 genes were down-regulated. **b**, Normalized enrichment score of differentially expressed pathways in the presence of phage clustered based on KEGG database at 12-hour sampling time point (GSEA, *FDR* < 0.05, *n* = 5). Pathways associated with flagellar assembly, bacterial chemotaxis and two-component signal transduction system were down-regulated, while type IV secretion system was significantly up-regulated in the presence of phage.

**Supplementary Figure 4**

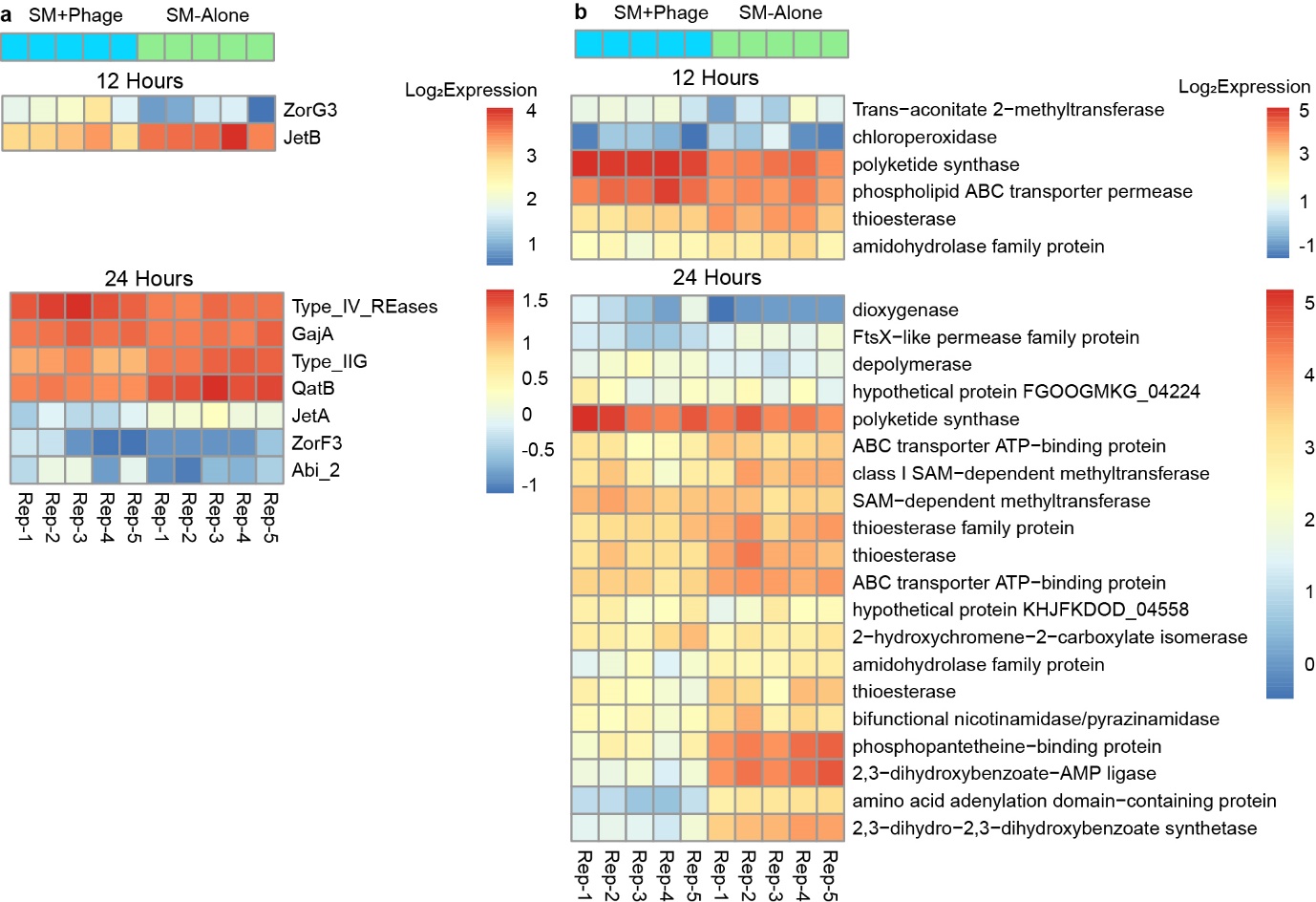

**Supplementary Figure 4.** **Differential expression of *S. maltophilia* genes related to potential phage resistance (a) and secondary metabolism (b) in the absence and presence of phage.** Read counts were normalized by transcripts per million (TPM) method and shown as log transformed data. Only genes that were significantly differentially expressed (DEseq, *P* < 0.05) are included in heatmaps.

**Supplementary Figure 5**

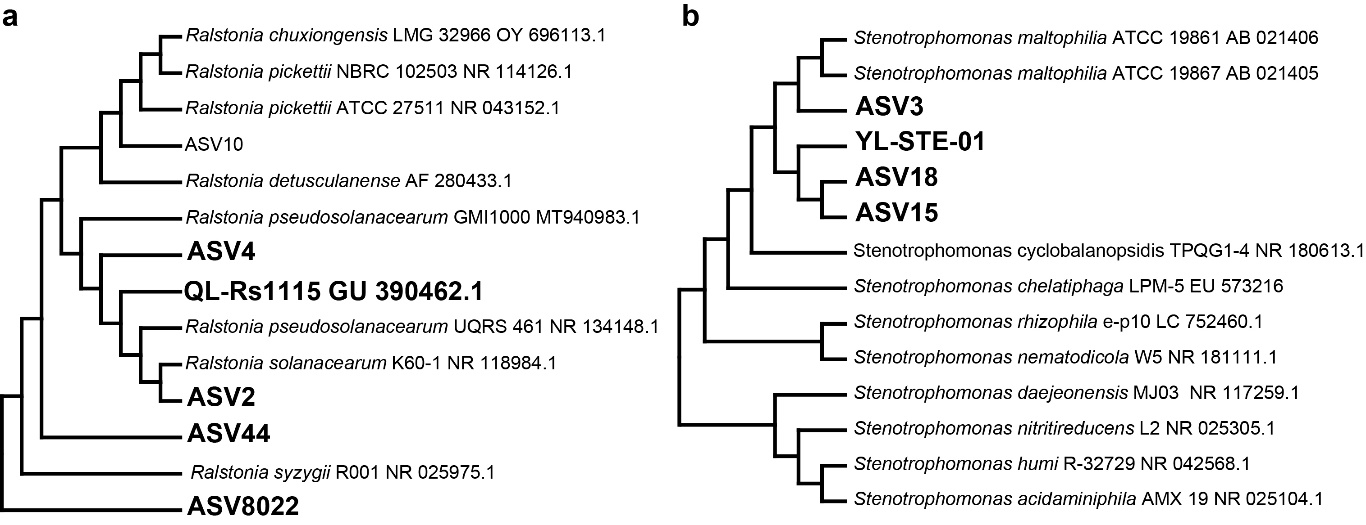

**Supplementary Figure 5. The taxonomic classification of ASVs belonging to *Ralstonia* (a) and *Stenotrophomonas* genera (b) and the relative abundance of *Ralstonia solanacearum* Species complex (RSSC) ASVs between treatments (b).** Phylogenetic tree was constructed based on 16s rRNA sequences using neighbor-joining method.

**Supplementary Figure 6**

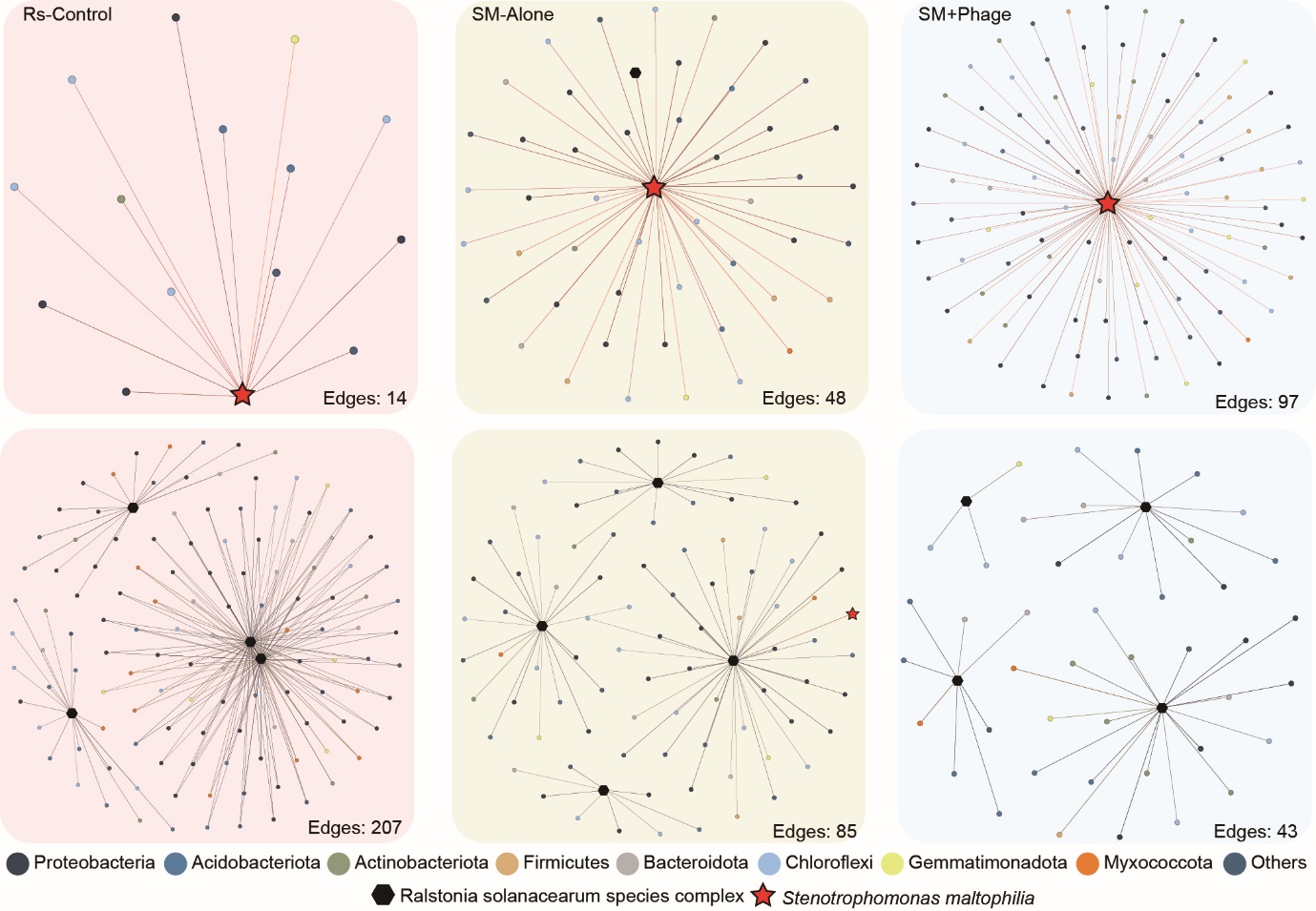

**Supplementary Figure 6. Sub-networks of ASVs of resident microbiota connected to ASVs belonging to *S. maltophilia* (upper) and RSSC (down) based on Fig. 4f.** The edges of each network are given below the networks. ASVs belonging to RSSC and *S. maltophilia* are highlighted with black hexagons and red stars, respectively.

**Supplementary Figure 7**

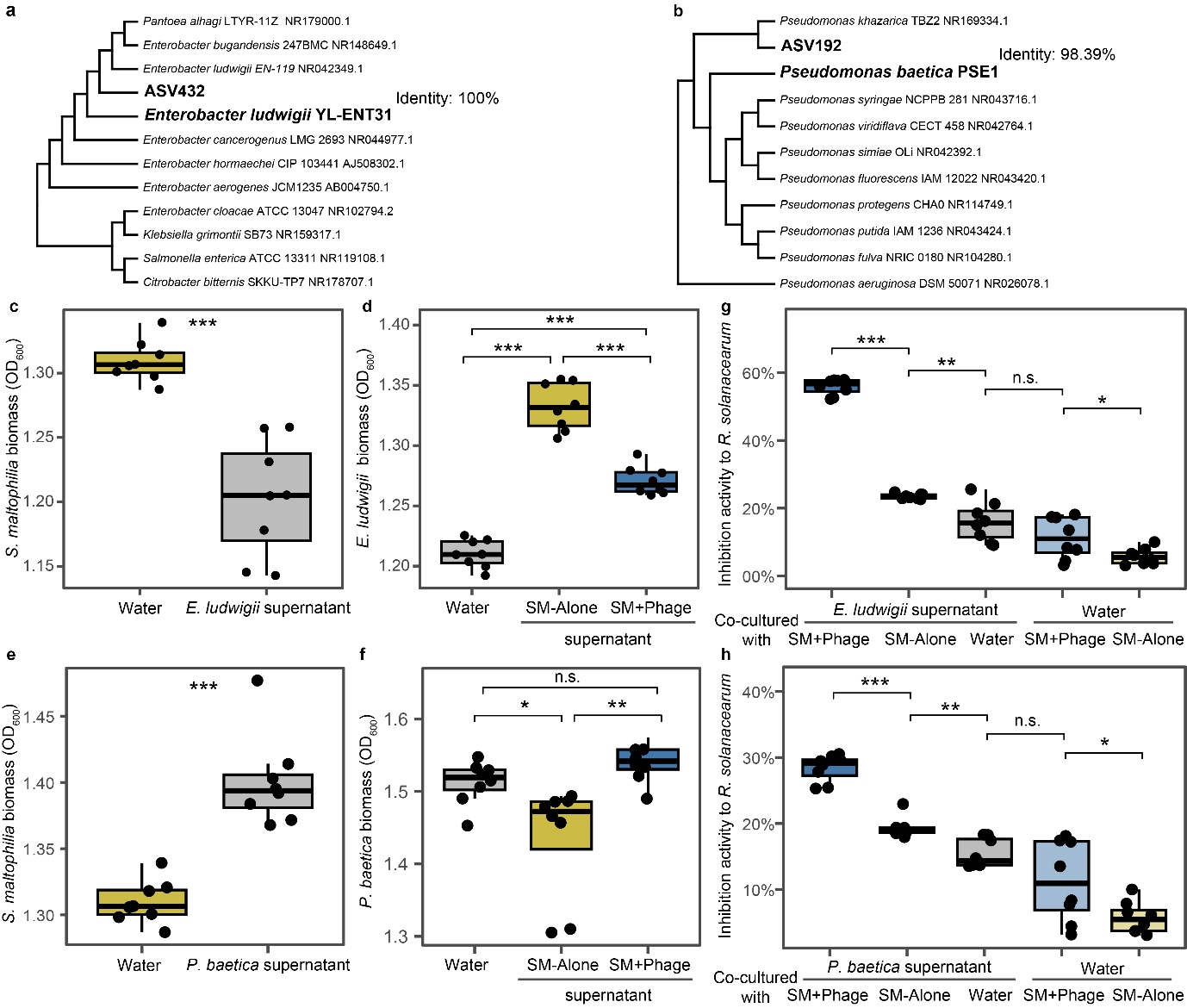

**Supplementary Figure 7.** **The effect of *S. maltophilia* phage on the interaction between *S. maltophilia* and *E. ludwigii* and *S. maltophilia* and *P. baetica* and consequences for the inhibition of *R. solanacearum* growth. a-b**, Close taxonomic associations of ASV432 and *E. ludwigii* YL-ENT-31 (a) and ASV192 and *P. baetica* PSE1 (b) based on 16S rRNA sequence phylogenetic tree using neighbor-joining method. **c and e**, *S. maltophilia* biomass when treated with water or bacterial supernatant of *E. ludwigii* and (c) *P. baetica* (e). **d and f**, *E. ludwigii* (d) and *P. baetica* (f) biomass treated with water and *S. maltophilia* supernatant cultured in the absence and presence of phage. **g-h**, Inhibition activity to *R. solanacearum* of *E. ludwigii* (g) or *P. baetica* (h) supernatant cultured with water and *S. maltophilia* supernatant previously cultured in the absence and presence of phage. Water added with same amount as *S. maltophilia* supernatant previously cultured in the absence and presence of phage was used as control. Unpaired t-test, n.s.: *P* > 0.05, **P* < 0.05, ***P* < 0.01, ****P* < 0.001, *n* = 8.

**Supplementary Figure 8**

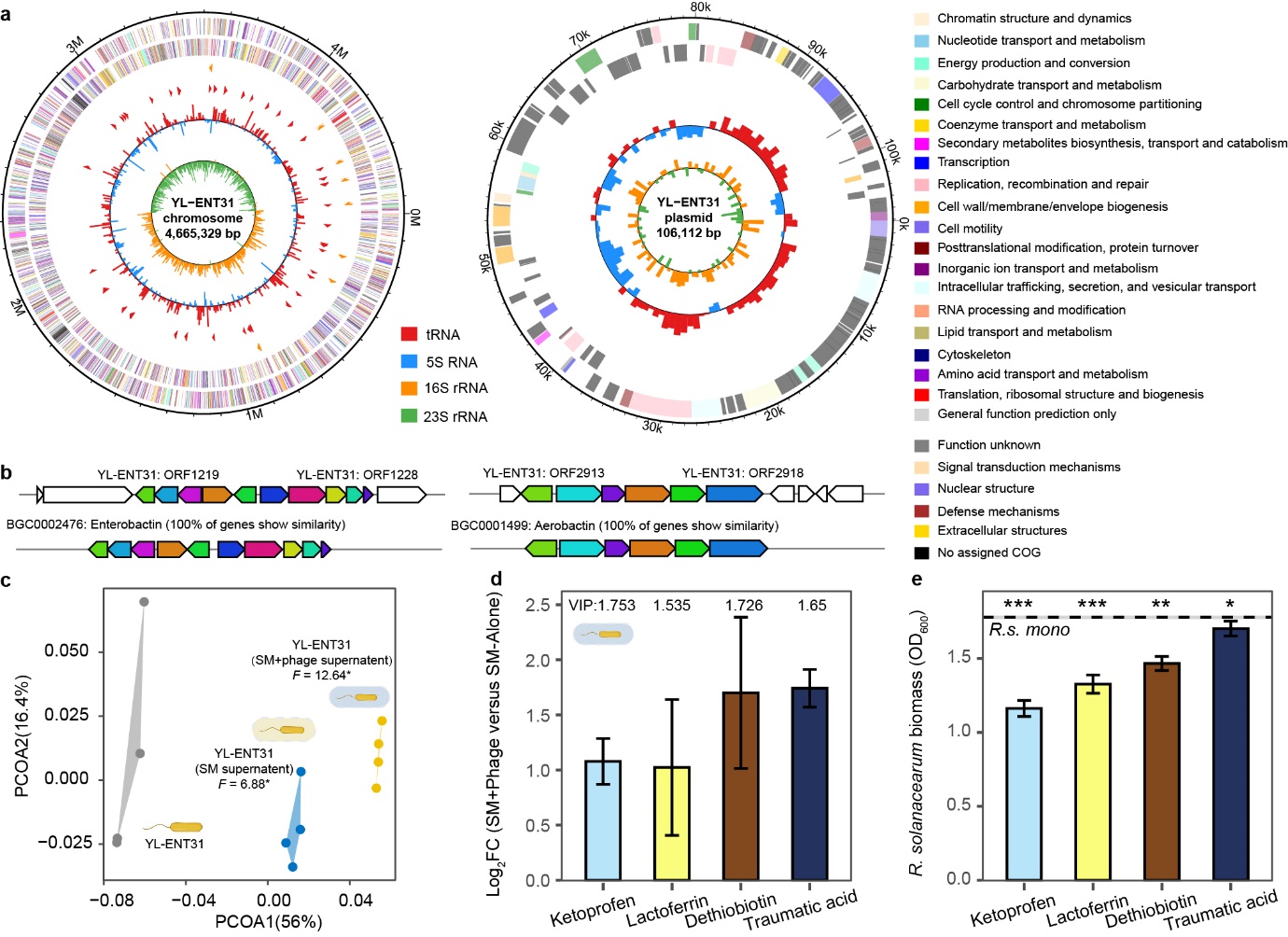

**Supplementary Figure 8. Effect of *S. maltophilia*-specific phage on the secondary-metabolite production by *E. ludwigii* YL-ENT-31*.* a**, The chromosomal (left) and plasmid (right) genome maps of *E. ludwigii* YL-ENT-31. **b**, Comparison of secondary metabolite biosynthesis related gene clusters based on antiSMASH. Gene clusters from ORF1219 to ORF1228 and ORF2913 to ORF2918 showed 100% similarity to gene clusters of Enterobactin and Aerobactin biosynthesis, respectively. **c**, Comparison of metabolomes of *E. ludwigii* supernatants when supplemented with *S. maltophilia*, which was co-cultured in the presence (SM+Phage) or absence (SM) of phage. (PERMANOVA, *: *P* < 0.05, *n* = 4). **d**, Fold change (mean ± SD) of ketoprofen, lactoferrin, dethiobiotin and traumatic acid abundance in the *E. ludwigii* supernatant metabolome affected by *S. maltophilia*-phage (log transformed, *P* < 0.05 by unpaired t-test for all compounds). The scores of variable important projection (VIP) are labeled above each metabolite. **e**, Reduction in *R. solanacearum* biomass (mean ± SD) treated with ketoprofen acid, lactoferrin, dethiobiotin and traumatic acid; dashed line with shading indicates the mean and standard deviation of *R. solanacearum* monoculture without any compounds (Unpaired t-test compared to mono culture control, *: *P* < 0.05, **: *P* < 0.01, ***: *P* < 0.001, *n* = 6).

**Supplementary Table 1**

Phage resistance and *R. solanacearum* inhibition activity of *S. maltophilia* mutants that had evolved in the presence or absence of phages (evolved alone).

| *S. maltophilia* treatment | Line | Phage  resistance | Inhibition activity  to *R. solanacearum* |
| --- | --- | --- | --- |
| Evolved with phage | 1 | 8.22% | 48.88% |
|  | 2 | 7.18% | 48.44% |
|  | 3 | 6.18% | 46.25% |
|  | 4 | 8.70% | 45.60% |
| Evolved alone | 1 | -40.34% | 30.69% |
|  | 2 | -34.53% | 25.12% |
|  | 3 | -38.68% | 24.47% |
|  | 4 | -35.62% | 29.90% |
| Ancestral *S. maltophilia* |  | -38.94% | 27.19% |

**Supplementary Table 2**

Predicted protein-protein interactively for ancestral/mutant *ssb* protei binding with Abi family protein, Type IV REases, *StmPr1* and polyketide synthase by PDBePISA^26^ and PRODIGY^27^.

| *ssb* protein source | Target protein | Interface area (Å^2^) | ΔG (kcal mol^-1^) | Intermolecular contacts |
| --- | --- | --- | --- | --- |
| Ancestral | Abi family protein | 3090.0 | -22.4 | 263 |
| Mutant |  | 3698.4 | -29.3 | 360 |
| Ancestral | Type IV REases | 3079.3 | -26.3 | 314 |
| Mutant |  | 3894.7 | -36.6 | 394 |
| Ancestral | *StmPr1* | 2948.2 | -21.7 | 294 |
| Mutant |  | 3732.5 | -32.8 | 358 |
| Ancestral | Polyketide synthase | 2430.6 | -17.6 | 209 |
| Mutant |  | 2318.2 | -19.9 | 216 |

**Supplementary Table 3**

Statistics for comparing tomato rhizosphere microbiome beta diversity between different treatments based on PERMANOVA analysis of Bray-Curtis distance.

| Treatment1 | Treatment2 | Df | Sum of Sqs | R^2^ | *F*-value | *P*-value |
| --- | --- | --- | --- | --- | --- | --- |
| Pre: SM+Phage | Pre: Control | 1 | 0.53901 | 0.44588 | 4.8279 | 0.031 |
|  |  | 6 | 0.66987 | 0.55412 |  |  |
|  |  | 7 | 1.20888 | 1 |  |  |
| Pre: SM | Pre: Control | 1 | 0.47149 | 0.36883 | 3.5062 | 0.027 |
|  |  | 6 | 0.80686 | 0.63117 |  |  |
|  |  | 7 | 1.27835 | 1 |  |  |
| Post: SM+Phage | Post: Rs-Control | 1 | 0.66168 | 0.44384 | 4.7883 | 0.024 |
|  |  | 6 | 0.82913 | 0.55616 |  |  |
|  |  | 7 | 1.49081 | 1 |  |  |
| Post: SM | Post: Rs-Control | 1 | 0.43469 | 0.31514 | 2.7609 | 0.03 |
|  |  | 6 | 0.94468 | 0.68486 |  |  |
|  |  | 7 | 1.37936 | 1 |  |  |
| Post: SM+Phage | Pre: Control | 1 | 0.17755 | 0.18788 | 1.388 | 0.076 |
|  |  | 6 | 0.76748 | 0.81212 |  |  |
|  |  | 7 | 0.94503 | 1 |  |  |
| Post: SM | Pre: Control | 1 | 0.24493 | 0.21714 | 1.6642 | 0.027 |
|  |  | 6 | 0.88303 | 0.78286 |  |  |
|  |  | 7 | 1.12796 | 1 |  |  |
| Post: Rs-Control | Pre: Control | 1 | 0.6459 | 0.42205 | 4.3815 | 0.034 |
|  |  | 6 | 0.8845 | 0.57795 |  |  |
|  |  | 7 | 1.5304 | 1 |  |  |
